# Rapamycin rescues loss-of-function in blood-brain barrier-interacting regulatory T cells

**DOI:** 10.1101/2022.10.21.513147

**Authors:** Paulien Baeten, Ibrahim Hamad, Cindy Hoeks, Michael Hiltensperger, Bart Van Wijmeersch, Veronica Popescu, Lilian Aly, Veerle Somers, Thomas Korn, Markus Kleinewietfeld, Niels Hellings, Bieke Broux

## Abstract

In many autoimmune diseases, FOXP3^+^ regulatory T cells (Tregs) skew towards a pro-inflammatory and non-suppressive phenotype and are therefore unable to control the exaggerated autoimmune response. This may largely impact the success of autologous Treg therapy which is currently under investigation for treatment of autoimmune diseases, including multiple sclerosis (MS). Thus, there is a need to ensure *in vivo* stability of Tregs before successful Treg therapy can be applied. Using a murine genetic fate-mapping model, we demonstrate that inflammatory exFOXP3 T cells accumulate in the central nervous system (CNS) during experimental autoimmune encephalomyelitis (EAE). In a human *in vitro* BBB model, we discovered that interaction with inflamed blood-brain barrier (BBB)-endothelial cells induces loss of suppressive function in Tregs. Transcriptome analysis further revealed that Tregs which migrated across inflamed BBB-endothelial cells *in vitro* have a pro-inflammatory Th1/17 signature and upregulate the mTORC1 signaling pathway compared to non-migrated Tregs. These findings suggest that interaction with BBB-endothelial cells is sufficient to affect Treg function, and that transmigration triggers an additive pro-inflammatory phenotype switch, which was also seen in CNS-derived exFOXP3 T cells of EAE mice. *In vitro* treatment of migrated human Tregs with the clinically-approved mTORC1 inhibitor rapamycin completely restored the loss of suppressive function. Finally, flow cytometric analysis indicated an enrichment of inflammatory, less suppressive CD49d^+^ Tregs in the cerebrospinal fluid of MS patients, thereby underscoring the relevance of our findings for human disease. In sum, our findings provide firm evidence that the inflamed BBB affects human Treg stability, which can be restored using a mTORC1 inhibitor. These insights can help in significantly improving the efficacy of autologous Treg therapy of MS.

## Introduction

Re-establishing tolerance is the holy grail of treating autoimmune diseases. More than a decade ago, researchers started to investigate the body’s own immunosuppressive machinery, resulting in clinical trials using regulatory T cell (Treg)-based cell therapies [1]. Although reported to be safe, efficacy is largely lacking [2, 3]. It is suggested that this failure could be due to the described loss of Treg phenotype in autoimmunity [4-9], as we and others showed for multiple sclerosis (MS) [10-12]. MS is a chronic demyelinating disorder of the central nervous system (CNS), mediated by an autoimmune reaction. The current paradigm states that, after an initiating event (potentially Epstein-Barr virus infection [13]) in a genetically susceptible individual, peripherally activated autoreactive CD4^+^ T helper (Th) cells initiate blood-brain barrier (BBB) disruption, thereby causing influx of circulating inflammatory immune cells into the brain (reviewed in [14-16]). This exaggerated local immune response leads to demyelination and axonal loss in both the white and grey matter [17]. As a result, symptoms like loss of vision, memory and sensation and even paralysis can occur. Current treatments mainly focus on relieving inflammation and thus delaying disease progression, while therapies targeting demyelination and neurodegeneration are missing. Therefore, new therapies avoiding general immunosuppression and inducing remyelination and neuroregeneration are needed. Interestingly, forkhead box protein 3^+^ (FOXP3^+^) CD4^+^CD25^high^CD127^low^ Tregs are not only able to control the exaggerated immune response, but have also been shown to induce regeneration in the CNS via amphiregulin (AREG) and CCN3 [18-21]. Therefore, Treg-based therapies may provide a promising therapeutic option for MS.

In homeostatic conditions, the BBB limits access of soluble factors and immune cells from the blood into the brain. This specialized function is mediated by the expression of intercellular tight junctions and specific transporters on BBB-endothelial cells (BBB-ECs), as well as the glia limitans produced by astrocytes [22]. In MS, the integrity of the BBB is affected and expression of chemokines and adhesion molecules is induced, thereby favouring immune cell entry into the perivascular space, and ultimately, the brain parenchyma [23]. Although Tregs intrinsically have an increased migratory capacity compared to effector T cells (Teff), this function is impaired in relapsing-remitting (RR)MS patients [24]. However, Tregs are found in the cerebrospinal fluid (CSF) of RRMS patients [11, 12, 25] and in active brain lesions [25]. In experimental autoimmune encephalomyelitis (EAE), Tregs found in the CNS are unable to suppress CNS-resident Teff. This effect is mediated by interleukin-6 (IL-6) and tumour necrosis factor-α (TNF-α) produced by these Teff [26]. In a different study, interferon-γ (IFN-γ)- producing Tregs with downregulated FOXP3 expression (exFOXP3 T cells) were found in the CNS of EAE animals [27]. In addition, it was shown that monocytes migrating through the BBB differentiate into dendritic cells which promote inflammatory Th17 cell polarisation [28]. Indeed, human Tregs are also known to skew towards pro-inflammatory Th1- and Th17-like cells in autoimmune diseases, depending on the micro-environment [29, 30]. Together, this raises the question whether Tregs are affected by BBB transmigration in a neuroinflammatory setting as well. In the context of Treg therapy, this *in vivo* Treg instability needs to be addressed to ensure safe and successful treatment [2, 3].

Here, we investigated the phenotype and function of Tregs interacting with an inflamed BBB. We demonstrate that inflammatory exFOXP3 T cells accumulate in the CNS of FOXP3^Cre-GFP^ Rosa^RFP^ fate-mapping mice during the course of EAE. In a human *in vitro* model of the BBB, we found that the suppressive capacity of Tregs interacting with inflamed BBB-ECs was affected. Further analysis on the transcriptome of migrated Tregs derived from both healthy donors (HD) and untreated RRMS (uRRMS) patients was performed using RNA sequencing (RNAseq). We show that inflammatory Th1/17 pathways and mammalian target of rapamycin complex 1 (mTORC1) signaling are upregulated in human BBB-transmigrated Tregs compared to non-migrated Tregs. In uRRMS-derived Tregs specifically, Th17-related pathways were increased after migration, suggesting a pre-existing susceptibility to Th17 skewing of Tregs in MS patients. In addition, downregulation of AREG in uRRMS-derived migrated Tregs suggests a disease-specific loss of regenerative capacity after BBB transmigration. Together, these results suggest that interaction with BBB-ECs is sufficient to affect Treg function, and that transmigration triggers an additive pro-inflammatory phenotype switch in Tregs, which is most pronounced in uRRMS patients. This finding was corroborated in EAE mice, where CNS-derived exFOXP3 T cells display similar changes. Treatment of human migrated Tregs with the mTORC1 inhibitor rapamycin restored and even augmented Treg function. Importantly, uRRMS-derived Tregs were still sensitive to rapamycin. Finally, an enrichment of inflammatory, less suppressive CD49d^+^ Tregs was found in the CSF of untreated RRMS patients, thereby underscoring the relevance of our findings for human disease. Altogether, our results show that interaction with inflamed BBB-ECs induces loss-of-function in Tregs, and that transmigration induces an additive pro-inflammatory skewing, which should be addressed before applying autologous Treg therapy to patients with autoimmune diseases such as MS.

## Results

### Loss of phenotype and function in BBB-transmigrated Tregs

Studies in mice have shown that the CNS is enriched for Tregs that are pro-inflammatory and non-suppressive [26, 27]. Since it was previously shown that migration across the BBB affects the phenotype of monocytes [28], we investigated whether the BBB is involved in the reported Treg phenotype switch and loss of function. To interrogate the FOXP3 expression of Tregs after migration into the CNS, we induced active EAE by myelin oligodendrocyte glycoprotein (MOG_35-55_) immunization in FOXP3^Cre-GFP^ Rosa^RFP^ fate-mapping mice. This transgenic model enables discrimination of bona fide FOXP3^+^ Tregs (RFP^+^GFP^+^) from exFOXP3 T cells (RFP^+^GFP^-^) in different tissues along the course of the disease using flow cytometry (Fig. 1A-E and Fig. S1). Interestingly, we found that exFOXP3 T cells accumulate in the CNS over time (Fig. 1D). In the chronic phase of the disease, exFOXP3 T cells are significantly more present in the CNS compared to the periphery (Fig. 1E). In addition, FOXP3^+^ Tregs outnumber the proportion of exFOXP3 T cells in lymph nodes (LN), while both cell populations are found in equal proportions in the CNS. Altogether, these results suggest that exFOXP3 T cells are enriched in the inflamed CNS *in vivo*.

**Fig. 1.**
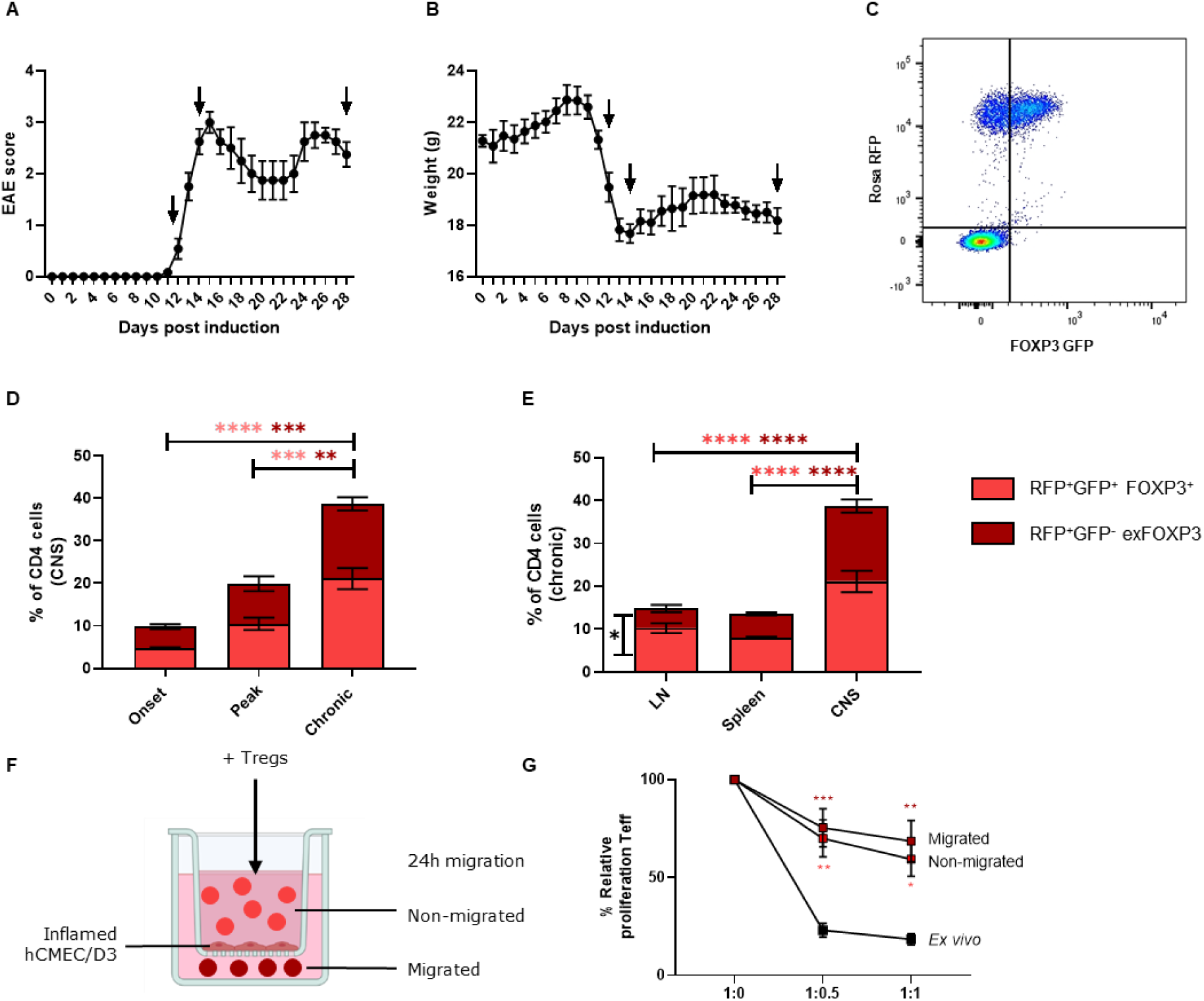
Lost phenotype and function in BBB-migrated Tregs. **A-E**. EAE was induced in female FOXP3^Cre-GFP^ Rosa^RFP^ fate-mapping mice. Tissues were collected at different time points (represented by the arrows) and immune cells were isolated from spleen, lymph nodes or pooled brain and spinal cord (CNS). **A**. Course of EAE with onset (12 dpi), peak (14 dpi) and chronic phase (28 dpi). n=4-12. **B**. Weight loss correlates to the disease course. n=4-12. **C**. Representative dot plot of RFP vs. GFP in splenocytes. Gating strategy in Fig. S1. **D**. Percentage of FOXP3^+^ Tregs and exFOXP3 T cells in the CNS along the EAE course. n=3-5; 2way ANOVA with Bonferroni’s multiple comparisons test. **E**. Percentage of FOXP3^+^ Tregs and exFOXP3 T cells at the chronic phase. n=4; 2way ANOVA with Bonferroni’s multiple comparisons test. Data are plotted as mean ± SEM. **F-G**. Suppressive capacity of fresh *ex vivo* isolated Tregs, *in vitro* non-migrated and migrated human HD-derived Tregs. **F**. Schematic representation of Boyden chamber migration assay. Tregs are FACS-sorted from PBMCs and loaded on the chamber. After 24h, migrated and non-migrated Tregs were collected for further analysis. **G**. Suppressive capacity of *ex vivo* Tregs, *in vitro* non-migrated or migrated human HD-derived Tregs. Percentage proliferation represents CellTrace dilution of Teff in suppression assays with different ratio’s of Teff and Tregs (given as Teff:Tregs). Relative proliferation is normalized to 1:0 condition (100%). n=4-8; 2way ANOVA with Bonferroni’s multiple comparisons test compared to *ex vivo* condition per ratio. Gating strategy in Fig. S2.*: p<0.05; **: p<0.01; ***: p<0.001; ****: p<0.0001.

To identify whether BBB-endothelial cells (ECs) are responsible for Treg instability, we used the human brain endothelial cell line hCMEC/D3 in a modified Boyden chamber migration assay to mimic the inflamed MS BBB *in vitro* (Fig. 1F) [31]. This model enables us to look at the direct effect of BBB-ECs on Treg function. Therefore, Tregs were allowed to migrate across inflamed BBB-ECs, after which the suppressive capacity of both migrated and non-migrated Tregs was determined. Compared to *ex vivo* Tregs, both migrated and non-migrated Tregs displayed a reduced capacity to suppress Teff proliferation (Fig. 1G), suggesting that interaction with inflamed BBB-ECs is sufficient to induce a loss-of-function in Tregs.

### BBB transmigration activates inflammatory and mTORC1 pathways in human and mouse Tregs

To identify the pathways affected in BBB-transmigrated human Tregs, RNAseq was performed on untouched, migrated and non-migrated Tregs from 5 HD and 4 untreated RRMS (uRRMS) patients (Fig. 2 and Fig. 3). Clustering analysis of these samples shows that untouched Tregs from HD and uRRMS cluster separately from migrated and non-migrated Tregs (Fig. 2A). The transcriptomic profile of non-migrated Tregs (those that are in contact with BBB-ECs but have not migrated through), significantly differs from untouched Tregs, indicating that interaction with inflamed BBB-ECs is sufficient to induce significant transcriptional changes in Tregs. Furthermore, clustering analysis indicates additional transcriptional changes in migrated Tregs, compared to non-migrated Tregs. Together with the *in vivo* data showing accumulation of exFOXP3 T cells in the CNS, this highlights the importance of identifying differences between migrating and non-migrating (but nonetheless interacting) Tregs.

**Fig. 2.**
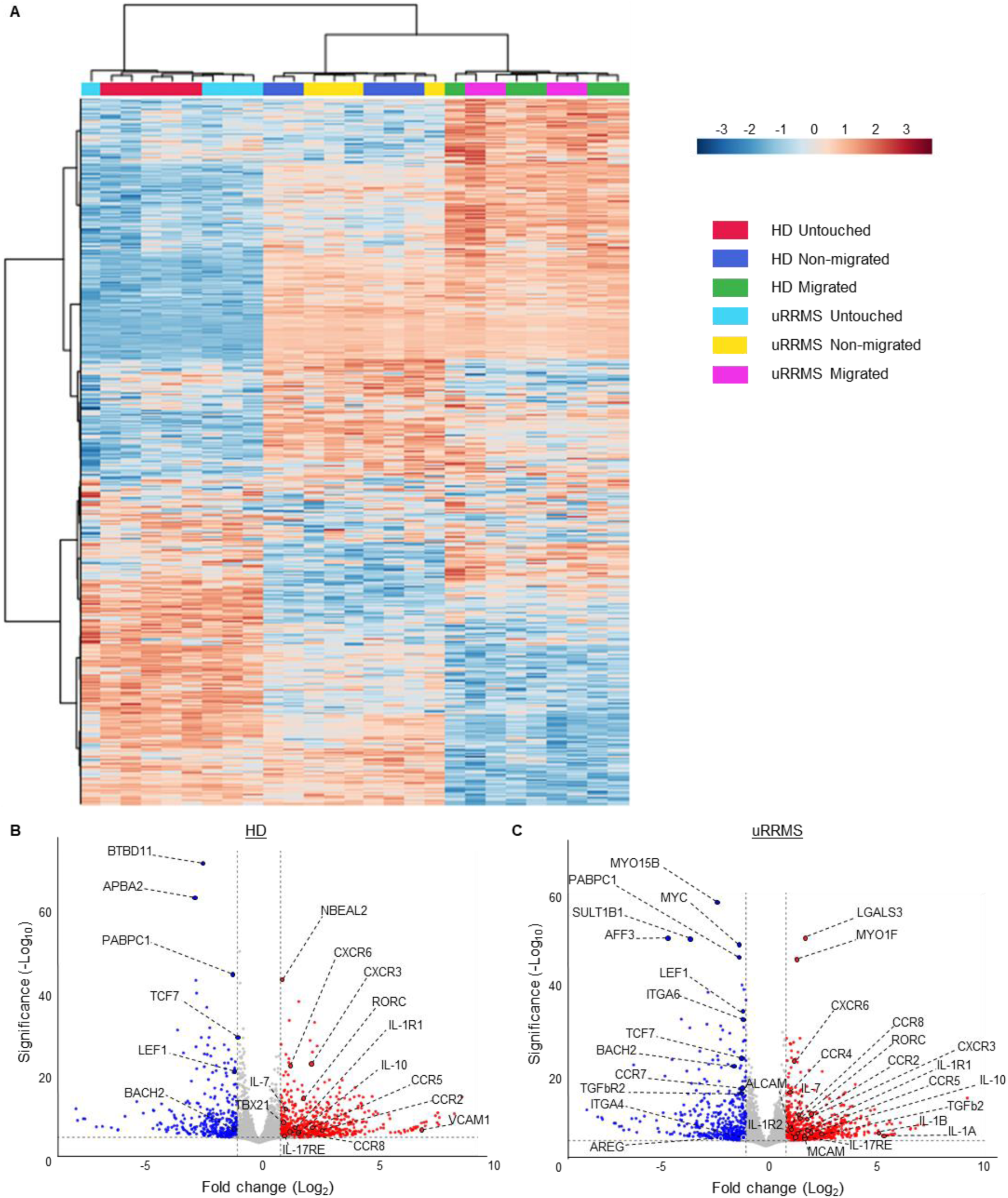
Human BBB-EC-transmigrated Tregs present a dysfunctional, pro-inflammatory phenotype. HD and uRRMS-derived Tregs were loaded on a Boyden chamber migration assay as represented in Fig. 1F. After 24h, untouched, migrated and non-migrated Tregs were collected and bulk RNAseq was performed. **A**. Hierarchical clustering showing changes in gene expression of the experimental conditions. Relative gene expression is indicated by colour: upregulation in red and downregulation in blue. Genes and samples with similar expression were automatically grouped (left and top trees). Expression values are shown as z-scores. Benjamini-Hochberg adjusted p-value (FDR <0.05) was used to determine DEGs. **B-C**. Volcano plot of DEGs in migrated vs. non-migrated Tregs for HD (B) and uRRMS (C). X-axis shows the log2 fold change for ratio migrated/non-migrated. Y-axis shows statistical significance (FDR-adjusted p-value). Up- and downregulated genes are coloured red or blue respectively when adjusted p-value<0.05 and |log2FoldChange|>1. Inflammation-, migration-, and regulation-related genes are highlighted. n=5 (HD) and n=4 (uRRMS).

**Fig. 3.**
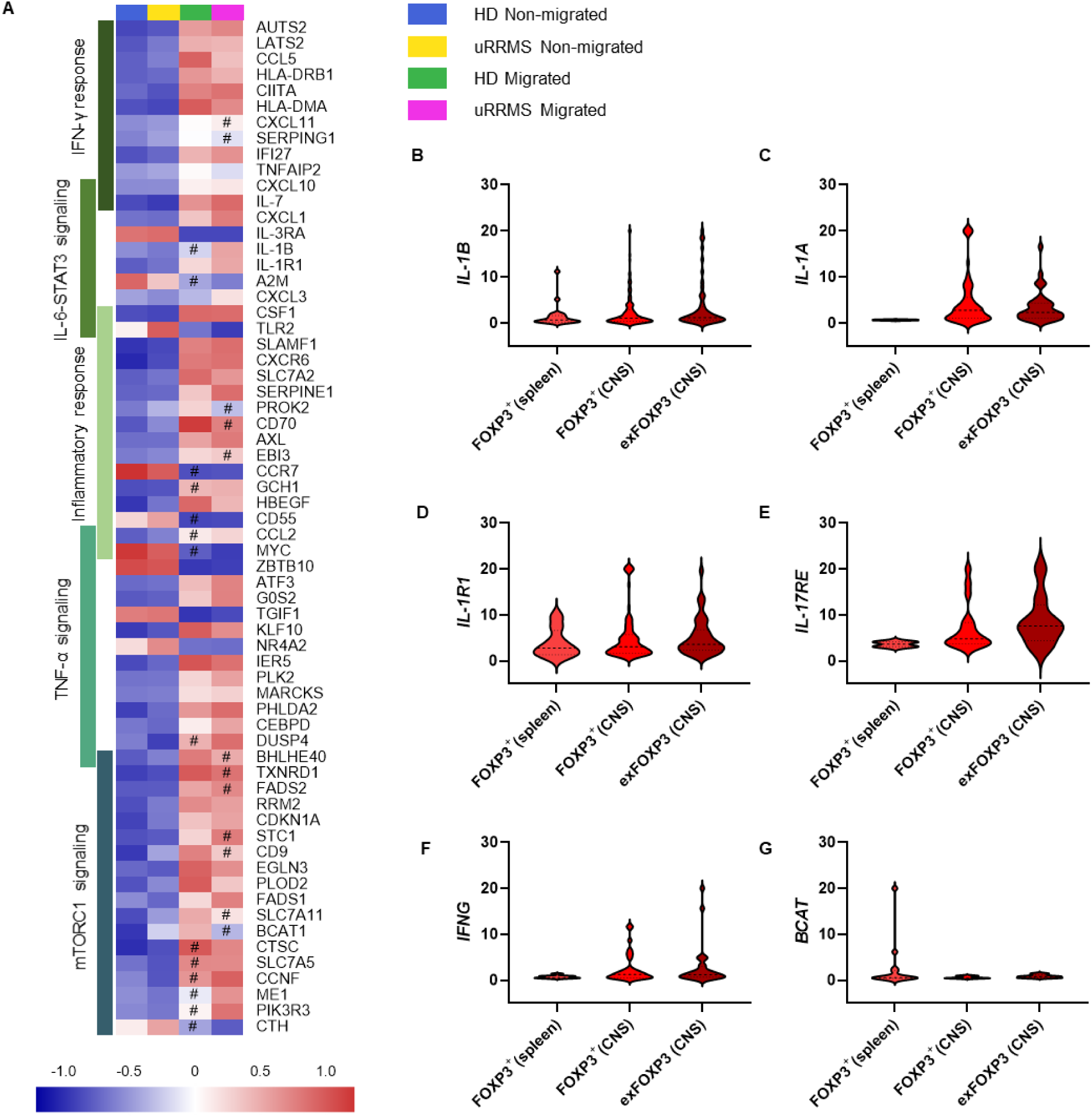
Pathway analysis of migrated Tregs shows upregulation of inflammatory phenotype and mTORC1 signaling. **A**. Differentially expressed genes (top 10 most significant) of the RNAseq experiment were grouped into following pathways: inflammatory response, TNF-α signaling, mTORC1 signaling, IFN-γ response, and IL-6-STAT3 signaling (only enriched in migrated Tregs derived from uRRMS patients). Pathways were identified using GSEA (Table S2 and Fig. S3). Relative gene expression is indicated by colour: upregulation in red and downregulation in blue. n=5 (HD) and n=4 (uRRMS); expression values are given as z-scores; Benjamini-Hochberg adjusted p-value (FDR <0.05) and|log2FoldChange|>1; non-significant differences (FDR>0.05 or |log2 FoldChange|>1) are marked with #. **B-G**. EAE was induced in female FOXP3^Cre-GFP^ Rosa^RFP^ fate-mapping mice. Tissues were collected at peak (18 dpi, Fig. S1) and RFP^+^GFP^+^ FOXP3^+^ Tregs and RFP^+^GFP^-^ exFOXP3 T cells were FACS-sorted from spleen and CNS (brain and spinal cord pooled). Gating strategy in Fig. S1. Single-cell RNAseq was performed and selected DEGs in CNS-derived RFP^+^GFP^-^ exFOXP3 T cells, CNS- and spleen-derived RFP^+^GFP^+^FOXP3^+^ Tregs are shown.

To identify specific transcriptional changes related to BBB transmigration, significant differences in gene expression between non-migrated and migrated Tregs in both HD and uRRMS patients were analysed and illustrated in volcano plots (Fig. 2B-C). Here, we found an increased expression of inflammation-related (e.g. *TNFRSF8/9, TBX21, RORC, IL-17RE, IL-1, IL-1R*) and migration-related (e.g. *VCAM1, ALCAM, MCAM, CXCR3, CCR4, CXCR6, CCR5)* differentially expressed genes (DEGs), while Treg phenotype-related genes (e.g. *TGFbR2, LEF1, TCF7, BACH2)* were downregulated. Of importance, *AREG* was downregulated only in MS-derived migrated Tregs, suggesting loss of regenerative capacity in these cells (Fig. 2C).

Next, ingenuity pathway analysis (IPA) (top 20 pathways in Table S1 and most applicable pathways in Fig. S3) and gene set enrichment analysis (GSEA) (top 20 pathways in Table S2 and most applicable pathways in Fig. S3) were performed to investigate relevant molecular pathways and regulatory networks that were affected in *in vitro* BBB-EC-transmigrated Tregs. As expected, actin cytoskeleton signaling pathway was increased (Fig. S3), which is related to the transmigration process. Moreover, several inflammation-related pathways were affected by BBB transmigration of Tregs, which are highlighted in Figure 3A. These include the (neuro-) inflammatory response, mTORC1 signaling, IFN-γ response (Th1 pathway) and TNF-α signaling for both HD and MS patients and IL-6-STAT3 signaling (IL-17 pathway) exclusively for MS patients (Fig. 3A, Fig. S3 and Table S1-2). In summary, we show that BBB-transmigrated Tregs display an additive pro-inflammatory phenotype switch on top of reduced suppressive function due to interaction with BBB-ECs, thereby resembling the Treg phenotype frequently observed in patients with autoimmunity [29, 30].

To validate our findings *in vivo*, we compared the gene expression of splenic RFP^+^GFP^+^ (FOXP3^+^) Tregs with RFP^+^GFP^+^ Tregs and RFP^+^GFP^-^ exFOXP3 T cells from the CNS of EAE mice at peak of disease by single-cell RNAseq (Fig. S1). Interestingly, the transcriptional pattern of murine CNS-derived exFOXP3 T cells is highly similar to that of human BBB-transmigrated Tregs, with comparable regulation of key genes like *IFNG, IL-1B, IL-1A, IL-1R1, IL-17RE* and *BCAT* (Fig. 3B-G). In particular, *BCAT* is a negative regulator of mTORC1 [32], which is decreased in CNS-isolated Tregs and exFOXP3 T cells compared to peripheral Tregs, suggesting that the mTORC1 pathway is upregulated similarly to human migrated Tregs. These data indicate that the pro-inflammatory skewing of BBB-transmigrating Tregs *in vitro*, combined with an unleashed mTORC1 pathway, also occur in an MS-like disease *in vivo*.

Next, flow cytometry and bead-based immunoassays were employed to validate selected targets and interesting pathways from the human RNAseq experiment at protein level. Chemokine receptors CCR6 and CCR4 were increased on migrated Tregs, in line with the identified migratory, inflammatory phenotype (Fig. 4A-B). In addition, the Th17-inducing molecule IL-6R [33] is more expressed on migrated Tregs, confirming pathway analysis of the RNAseq experiment (Fig. 4C). In line with this, we found that inflamed BBB-ECs produce high levels of IL-6 and IL-1β (Fig. S4), revealing a potential mechanism for Th17-skewing of Tregs. Next, using bead-based immunoassays, IFN-γ (Fig. 4D) and IL-10 (Fig. 4E) production was measured in the supernatants of suppression assays using migrated and non-migrated Tregs. Co-cultures of Teff with migrated Tregs resulted in higher IFN-γ levels in the supernatant compared to Teff co-cultured with non-migrated Tregs (Fig. 4D), suggesting an altered function of migrated Tregs. In line with our RNAseq data, IL-10 production is also higher in co-cultures when migrated Tregs are used, compared to Teff alone and also compared to co-culture with non-migrated Tregs (Fig. 4E). Although not sufficient to override the reduced suppressive function, this could indicate a compensation mechanism that is triggered in migrating Tregs. In summary, these data suggest that migration across BBB-ECs *in vitro* and *in vivo* induces an altered phenotype in Tregs, as highlighted by the upregulated pro-inflammatory Th1 and mTORC1 signaling pathways in migrated Tregs. In addition, key proteins are validated on migrated Tregs using flow cytometry. Interestingly, the Th17-related IL-6-STAT3 signaling pathway and the regenerative gene *AREG* are uniquely affected in MS-derived migrated Tregs. In addition, we confirmed this pro-inflammatory phenotype in CNS-isolated Tregs *in vivo*.

**Fig. 4.**
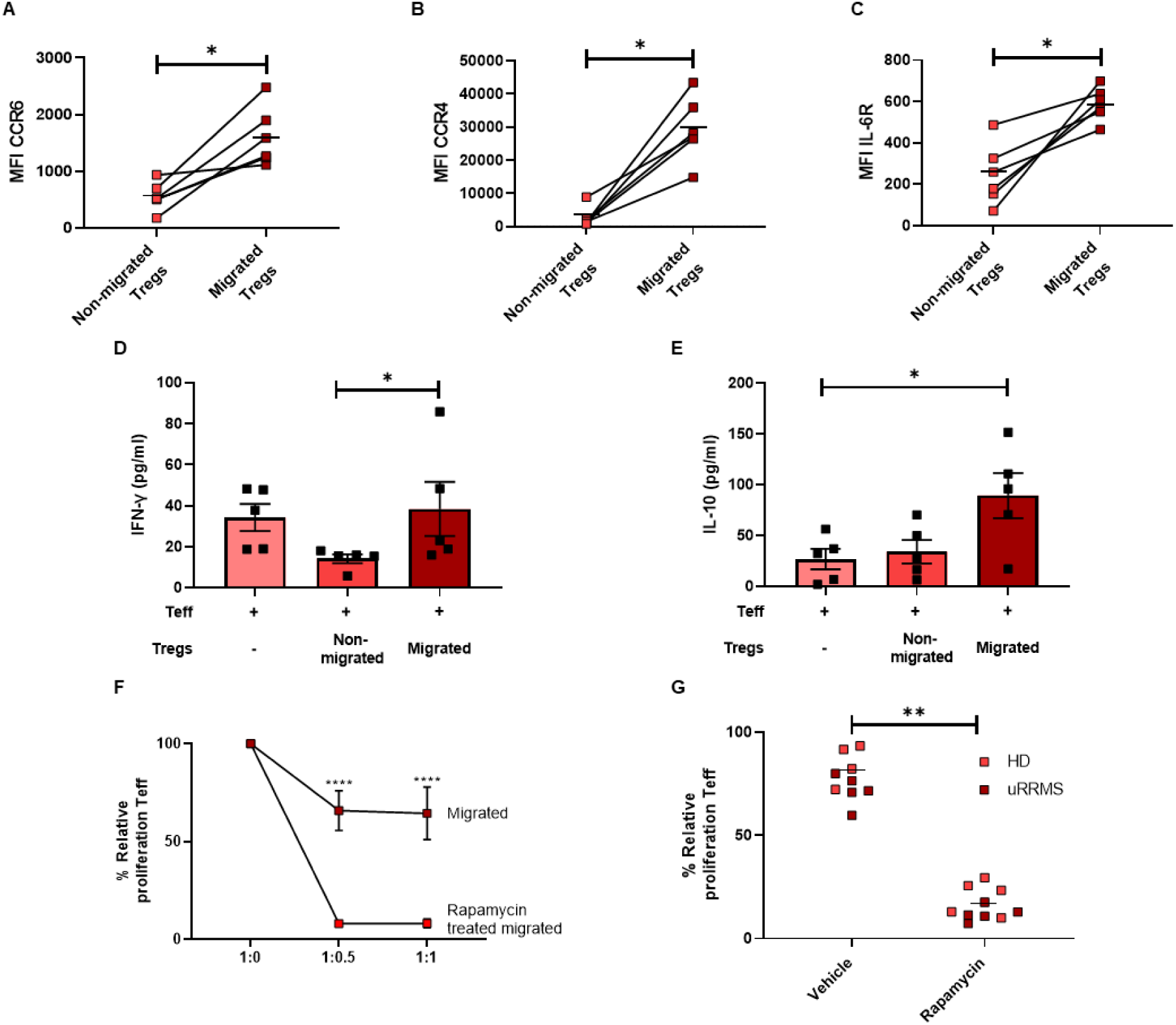
Pro-inflammatory phenotype of migrated human Tregs is restored by rapamycin. **A-C**. Using flow cytometry, molecules identified by pathway analysis were validated. MFI of CCR6 (**A**), CCR4 (**B**) or IL-6R (**C**) of live cells are shown. Horizontal bars represent group mean. n=6; Wilcoxon test. Gating in Fig. S8. **D-E**. Concentration of IFN-γ (**D**) and IL-10 (**E**) were measured in supernatants of Teff:Tregs co-cultures. n=5; Friedman test with Dunn’s multiple comparisons test. **F-G**. Tregs were treated with rapamycin (2μM) for 4h, washed and co-cultured with Teff in a suppression assay. **F**. Treatment of migrated HD-derived Tregs. Percentage proliferation represents CellTrace dilution of Teff. Cell ratio is given as Teff:Tregs. Relative proliferation is normalized to 1:0 condition (100%). Gating in Fig. S2. n=6; 2way ANOVA with Bonferroni’s multiple comparisons test. **G**. Frozen PBMC samples of HD and uRRMS patients were thawed and Tregs are isolated using FACS. Immediately after isolation, Tregs were treated with rapamycin (2μM) or vehicle (DMSO, 1/200) for 4h, washed and co-cultured with Teff in a suppression assay. Horizontal bars represent group mean. n=5 (HD), n=5(uRRMS); Wilcoxon test. *: p<0.05; **: p<0.01; ****: p<0.0001.

### Rapamycin treatment restores the suppressive capacity of migrated Tregs

One of the most prominently affected Treg function-related pathways by transmigration, is the mTORC1 signaling pathway. The mTORC1 inhibitor rapamycin is already being used in clinical trials (NCT01903473 and [34, 35]). To determine whether inhibition of mTORC1 restores Treg function after BBB transmigration, migrated Tregs were treated with rapamycin before co-culturing them with Teff in suppression assays. To prevent direct effects on Teff, Tregs were pre-incubated with rapamycin before co-culturing. Interestingly, rapamycin restored and even augmented the suppressive capacity of migrated Tregs (Fig. 4F; non-migrated Tregs in Fig. S5). In contrast, treating Tregs with rapamycin before BBB transmigration, did not result in a stable protection of Treg function (Fig. S5). To test whether Tregs from untreated uRRMS patients are still sensitive to rapamycin-enhanced suppressive capacity, Tregs were sorted from frozen PBMCs. Immediately after isolation, Tregs were treated with rapamycin or vehicle (DMSO) before co-culturing them with Teff in suppression assays. Indeed, these results show that resting Tregs of untreated uRRMS patients are equally sensitive to rapamycin as HD-derived Tregs (Fig. 4G). Together, our data demonstrate that rapamycin restores the suppressive function of migrated human Tregs, highlighting the importance of mTORC1 signaling for Treg stability in the context of MS.

### Pro-inflammatory and non-suppressive Tregs accumulate in CSF of MS patients

Having established that Tregs lose their phenotype and suppressive function after migration across inflamed BBB-ECs *in vitro* and in a mouse model of MS *in vivo*, we sought evidence to evaluate this phenotype switch in MS patients. Blood samples of 26 HD, 24 uRRMS patients, 25 first-line treated RRMS patients (flRRMS) and 23 untreated secondary-progressive MS patients (uSPMS) were analysed by flow cytometry. Pooled data were analyzed using FlowSOM, which uses self-organizing maps for visualization of flow cytometry data in an unbiased way [36]. We identified the presence of two Treg subpopulations in the peripheral blood, which are distinguished by CD49d expression (Fig. 5A-B, cluster 6 and 9). CD49d was previously found to be a marker for inflammatory and non-suppressive Tregs [37], although in this cohort, its expression is not different between blood samples of HD and MS patients (Fig S6). In contrast, analysis of paired blood and CSF samples of uRRMS patients at time of diagnosis, revealed that CD49d^+^ Tregs are significantly enriched in the CSF (Fig. 5C and Fig. S7). In line with our previous results, these data indicate that inflammatory and less-suppressive Tregs are not yet expanded in the blood of MS patients, however they do accumulate in the CSF.

**Fig. 5.**
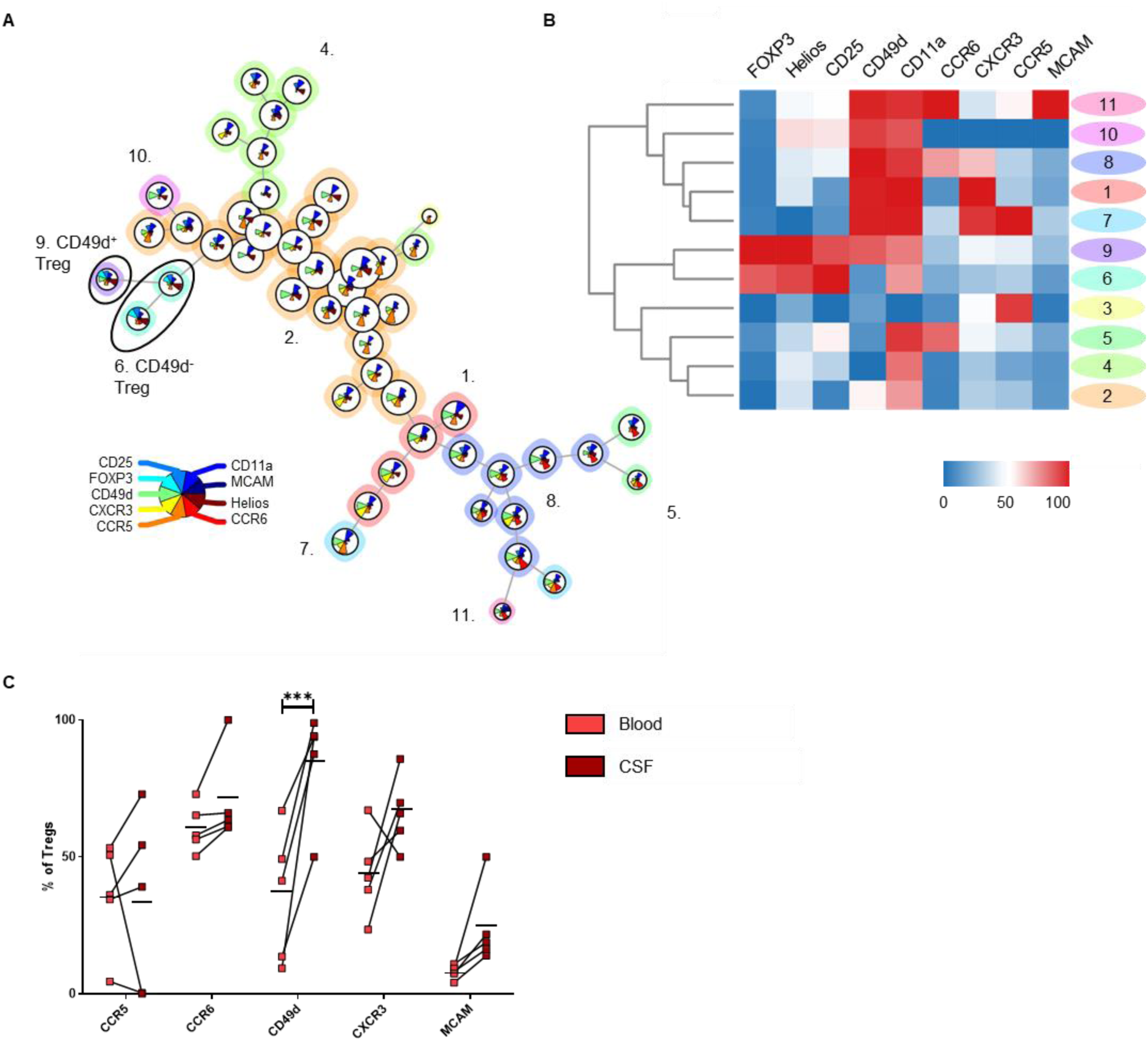
Pro-inflammatory CD49d^+^ Tregs accumulate in the CSF of MS patients. **A-B**. Frozen PBMC samples of HD and different MS patients were thawed and studied using flow cytometry. **A**. FlowSOM analysis on CD4 T cells identified two subpopulations of Tregs. **B**. Heatmap related to the FlowSOM analysis. n=26 (HD), n=24 (uRRMS), n=25 (flRRMS), n=23 (uSPMS) **C**. Fresh, paired blood and CSF samples were collected from uRRMS patients at diagnosis and studied using flow cytometry. Percentage expression of different migratory molecules within Tregs illustrated that CD49d^+^ Tregs are enriched in the CSF. Gating in Fig. S7. Horizontal bars represent mean of the group. n=5; 2way ANOVA with Bonferroni’s multiple comparisons test. ***: p<0.001.

## Discussion

The *in vivo* stability of Tregs is of crucial importance for Treg-based cell therapy in the context of autoimmunity [2, 3]. The identification of Treg instability at the site of inflammation, and its driving factors, will enable us to improve the efficacy and safety of Treg therapy. Loss of FOXP3 expression has been shown to occur in Tregs in inflammatory micro-environments, resulting in exFOXP3, non-suppressive cells [26, 27, 29, 38, 39]. Because of the autoreactive nature of Tregs [40], conversion to Teff poses a major risk for using Tregs as a cell-based therapy. Here, we investigated the role of Treg – BBB interaction and transmigration on the function and phenotype of Tregs in the context of MS. We found that, in a mouse model for MS, exFOXP3 cells accumulate in the CNS during the course of the disease. Using a human *in vitro* model, we provide evidence for a human Treg population with an altered phenotype and lost suppressive function after interaction with inflamed BBB-ECs. Even more, Tregs migrating across inflamed BBB-ECs display a pro-inflammatory phenotype which is most pronounced in uRRMS patients. Selective migration of a specific Treg subset cannot be ruled out, however, non-migrated Tregs, representing interacting yet non-EC-adherent cells without transmigration, represent a decreased suppressive capacity and a transcriptome already very divergent from untouched Tregs as well. Subsequently, transcriptomic analysis of migrated Tregs revealed that several inflammatory pathways (IFN-γ response, TNF-α signaling and IL-6-STAT3 pathway) and the mTORC1 signaling are upregulated compared to non-migrated Tregs. Although not sufficient to override the reduced suppressive function, migrated Tregs show increased IL-10 and *TGFb2* expression which could indicate a compensation mechanism. Key genes related to inflammation (*IFNG, IL-17RE, IL-1R1, IL1B and IL-1A*) and mTORC1 signaling (*BCAT*) were validated in spleen- and CNS-derived FOXP3^+^ Tregs and exFOXP3 T cells from EAE mice. Treatment of migrated Tregs with the mTORC1 inhibitor rapamycin fully restored their suppressive capacity. Of clinical relevance, Tregs from untreated uRRMS patients were still susceptible to rapamycin, which is important in a therapeutic context. Finally, we showed that less-suppressive and inflammatory Tregs, characterized by expression of CD49d, accumulated in the CSF of MS patients, supporting our *in vitro* human and murine *in vivo* results.

Previous studies identified non-suppressive and pro-inflammatory Tregs in the CNS of EAE mice, which thereby contribute to disease pathogenesis [26, 27]. These CNS-resident Tregs were previously shown to produce IFN-γ and were unable to suppress CNS-isolated Teff. Using a FOXP3^Cre-GFP^ Rosa^RFP^ fate-mapping mouse model, we now show that exFOXP3 T cells accumulate in the CNS along the EAE disease course. Using this model, one cannot rule out that this enrichment is not due to clonal expansion, however, due to low IL-2 levels in the CNS, this might be unlikely [41]. Further experiments will need to clarify these results. We demonstrate loss of Treg phenotype and function in human Tregs after contact with and migration across inflamed BBB-ECs *in vitro*, and we confirmed key inflammatory-related DEGs in CNS-derived exFOXP3 T cells and FOXP3^+^ Tregs *in vivo* in the fate mapping mouse model. These data suggest that although FOXP3^+^ Tregs are present in the CNS during EAE, they still represent a pro-inflammatory phenotype correlated with mTORC1 signaling.

The BBB consists of multiple cellular players, but we show in our *in vitro* unicellular model that BBB-ECs are sufficient to induce significant changes in migrating Tregs. Indeed, this is in line with the report of Ifergan *et al*. [28], who showed that migration of monocytes across BBB-ECs induced skewing towards Th17-differentiating dendritic cells. Even more, contact alone is sufficient to decrease the suppressive capacity of Tregs and induce transcriptomic changes, which are further aggravated by transmigration, suggesting a true alteration and no selective expansion. The conversion of migrating immune cells is suggested to be driven solely by factors expressed or produced by BBB-ECs, like IL-6 [42]. We confirmed production of IL-6 and IL-1β by inflamed hCMEC/D3 cells, while we showed that IL-6R and *IL-1R* expression are increased on migrated Tregs. However, no functional correlations were yet determined and further research is necessary. Indeed, literature has shown IL-6R expression on human Tregs as well [43], which renders Tregs, and more specifically thymic Tregs [44], more susceptible to Th17 skewing [45, 46]. Previous reports showed that IL-1 and IL-6 alone and together downregulate FOXP3 while inducing IL-17 expression on Tregs in a STAT3-dependent way [33, 47-51], also in EAE and rheumatoid arthritis [26, 29, 52]. As these cytokines are not BBB specific, these effects will not be limited to MS. Indeed, fibroblast-produced IL-6 also drives conversion of Tregs into IL-17-producing exFOXP3 cells accumulating in the inflamed joints of mice with autoimmune arthritis [29]. Tocilizumab, an IL-6R blocking antibody, is already approved for treatment of rheumatoid arthritis [53, 54] and has been successfully applied to hospitalized COVID-19 patients [55], showing its potential in dampening hyperinflammatory responses. In addition, blocking IL-1R in human Tregs was shown to prevent IL-17 production *in vitro* [48]. The marketed drug IL-1R antagonist Anakinra is currently under investigation in clinical trials for MS [56] (NCT04025554). These findings highlight the importance of the STAT3 signaling pathway in Th17 differentiation, and we here identified an MS-specific upregulation of this pathway in migrated Tregs. Further studies are needed to evaluate the effect of IL-6 and/or IL-1β blocking antibodies in preventing the detrimental inflammatory effects of BBB-ECs on migrating Tregs.

Kim *et al*. [49] showed that, in the context of hepatitis B liver failure, the STAT3-based Th17:Treg balance can be regulated by the mTORC1 inhibitor rapamycin. Rapamycin is shown to alleviate EAE symptoms and related Th1- and Th17-mediated inflammation while boosting Treg numbers [57, 58]. It is already being used in a clinical trials to treat MS patients (NCT01903473, [34, 35]) and to selectively expand highly suppressive Tregs *in vitro* for Treg cell therapy [2, 3]. Rapamycin was found to reduce contamination of Teff in Treg cultures, to maintain Treg function and phenotype and to prevent a pathogenic switch under Th skewing conditions [59-62]. Our *in vitro* RNAseq results show that the mTORC1 signaling is activated in migrated Tregs. Even more, expression of *BCAT*, a negative regulator of mTORC1, is decreased in CNS-derived Tregs *in vivo* [32]. Altogether, this prompted us to use rapamycin in our experiments, with the goal of rescuing the altered function of migrated Tregs. Indeed, we found that treatment of migrated Tregs with rapamycin recovered and even improved their suppressive capacity. Even more, Tregs from uRRMS patients were still susceptible to rapamycin, which is important in a therapeutic context. This reveals the potential of targeting any of the identified mechanisms in this set-up to stabilize Treg phenotype in the context of Treg-based cell therapy for autoimmune diseases. However, treatment of Tregs with rapamycin before BBB transmigration does not prevent loss-of-function. In the context of Treg therapy, this suggests that a transient treatment of *ex vivo* Tregs before infusion would not be sufficient to maintain *in vivo* stability, which is crucial for effective Treg therapy in MS.

When using MS-derived Tregs in the migration assay, we found unique upregulation of the Th17 signaling pathway, as well as downregulation of *AREG*. Tregs have been reported to induce regeneration in the CNS, using AREG [20]. This suggests that MS-derived migrated Tregs lose the ability to induce regeneration in the CNS due to loss of *AREG* expression. However, the regenerative capacity of MS-derived Tregs has not been studied yet, although this is of high interest since regenerative therapies are currently lacking for MS patients.

Altogether, these results indicate that interaction with BBB-ECs and transmigration is detrimental for human Tregs *in vitro* and *in vivo*. Interaction with BBB-ECs is sufficient to affect Treg function, and transmigration triggers an additive pro-inflammatory phenotype switch which is most pronounced in uRRMS patients. Interestingly, CNS-derived exFOXP3 T cells of EAE mice were transcriptionally highly similar to human BBB-transmigrating Tregs. Finally, we found that CD49d^+^ Tregs accumulate in the CSF of recently diagnosed uRRMS patients, thereby confirming our *in vitro* and murine results. These CD49d^+^ Tregs were previously reported to be inflammatory and less suppressive [37]. However, so far we cannot rule out that this effect does not result from selective migration of CD49^+^ Tregs. Altogether, the relevance of our findings was validated in the first clinical trial using Treg therapy in MS. Chwojnicki *et al*. showed that intrathecally administrated Tregs show more clinical potential compared to intravenously administered Tregs [64]. This finding supports our results and could explain disease progression in MS patients receiving intravenous infusion of Tregs. However, due to low patient numbers in this phase I trial, these conclusions should be taken with care. In order to move forward with Treg-based cell therapies for autoimmunity, overcoming and even preventing the observed phenotype switch will likely result in a stable and safe cellular product for clinical use. Our research identifies several pathways which can be used to restore Treg function, as we show with rapamycin, to ensure Treg stability, even in highly inflammatory conditions like the MS brain environment.

## Material & Methods

### Study design

This study investigated the role of Treg – BBB interaction on the function and phenotype of Tregs in the context of MS. To this end, EAE was induced in Treg fate-mapping mice in order to track FOXP3 Tregs and exFOXP3 T cells in spleen and CNS by flow cytometry and single-cell RNAseq. In addition, using hCMEC/D3 BBB-endothelial cells in a human *in vitro* model, human Tregs were analysed during the transmigration process. Afterwards, Tregs derived from MS patients and healthy donors were studied by RNAseq, flow cytometry and *in vitro* suppression assays, to determine Treg function and to identify underlying molecular mechanisms. Next, we aimed to restore the lost suppressive capacity of migrated Tregs by the means of rapamycin treatment *in vitro*. Finally, we aimed to validate these murine and *in vitro* human findings in MS patients. This was evaluated by flow cytometry using peripheral blood mononuclear cells of healthy donors and MS patients with different disease types and treatments, and paired blood-CSF samples of early RRMS patients.

### Animals

BAC-FOXP3^Cre-GFP^ mice [65-67] and Rosa26^fl^STOP^fl^RFP mice [68], with a C57BL/6 background, were crossed to obtain BAC-FOXP3^Cre-GFP^ Rosa26^fl^STOP^fl^RFP (FOXP3^Cre-GFP^ Rosa^RFP^) fate-mapping mice, enabling the discrimination of RFP^+^GFP^+^ FOXP3^+^ Tregs and RFP^+^GFP^-^ exFOXP3 T cells *in vivo*. Animals were housed in an accredited conventional animal facility under a 12h light/dark cycle and had free access to food and water. All mouse procedures were in accordance with the EU directive 2010/63/EU and were approved by the Hasselt University Ethics Committee for Animal Experiments.

### EAE induction

Ten week old female mice were subcutaneously injected with MOG_35-55_ emulsified in complete freund’s adjuvant (CFA) containing Mycobacterium tuberculosis, according to manufacturer’s instructions (Hooke Laboratories, Lawrence, USA). Immediately after immunization and on day 1, mice were intraperitoneally (i.p.) injected with 100ng/100μl pertussis toxin (PTX) to induce EAE. All animals were weighed daily and neurological deficits were evaluated using a standard 5-point scale (0: no symptoms; 1: limp tail; 2: weakness of hind legs; 3: complete paralysis of hind legs; 4: complete hind and partial front leg paralysis; 5: death). Analysis of the clinical EAE scores was performed using pooled data from three independent experiments (naïve, n=5; onset, n=4; peak, n=4; chronic, n=4). At EAE onset (12 days post induction (dpi)), EAE peak (14/18 dpi) and the chronic phase of disease (28 dpi) lymph nodes, spleen, spinal cord and brains were isolated after transcardial perfusion with Ringer’s solution. A single cell suspension from lymph nodes and spleen was derived by mechanical transfer through a 70μm cell strainer (Greiner Bio-One, Vilvoorde, Belgium). For the CNS, both enzymatic digestion, using collagenase D (Roche Diagnostics GmbH, Mannheim, Germany) and DNase I (Roche Diagnostics GmbH) and mechanical dissociation was performed, followed by a Percoll gradient (GE Healthcare, Diegem, Belgium). Isolated cells were stained with Zombie NIR and CD4 Pacific Blue (both Biolegend, San Diego, USA) and were acquired on BD LSRFortessa (BD Biosciences, Everbodegem, Belgium) and analysed using FlowJo 10.8.0 (BD Biosciences). In addition, cells isolated from spleen and CNS were stained with Zombie NIR and sorted using the FACSAria Fusion (BD Biosciences) for RFP^+^GFP^+^ FOXP3^+^ Tregs and RFP^+^GFP^-^ exFOXP3 T cells. The sorting strategy is depicted in Fig. S1 and purity of all sorts was confirmed to be >90%.

### Single-cell RNAseq

RFP^+^GFP^+^ FOXP3^+^ Tregs and RFP^+^GFP^-^ exFOXP3 T cells isolated from spleen and CNS from EAE mice were stained with sample tag antibodies (Ms Single Cell Sampling Multiplexing Kit, BD Biosciences). Next, cells were pooled and immediately used for single-cell capture using the BD Rhapsody Express system (BD Biosciences). Directly following cell capture, DNA library preparation was performed using the BD Rhapsody WTA Amplification Kit (BD Biosciences) according to manufacturer’s instructions. In short, after single-cell capture in the nanowell plate, Cell Capture beads were loaded and cells were lysed. Free floating mRNA was allowed to bind to the Cell Capture bead present in the same nanowell, and beads were collected. Next, reverse transcription and exonuclease I treatment were performed. Prior to mRNA amplification, Sample Tag products were denatured off the Cell Capture beads. WTA and Sample Tag libraries were prepared in parallel. After each amplification round and following index-PCR, the PCR products were purified using Agencourt AMPure XP beads (Beckman Coulter Diagnostics, Brea, USA). Concentration of final PCR products were determined using dsDNA High Sensitivity assay kit (Thermo Fisher Scientific, Waltham, USA) on the Qubit analyser (Thermo Fisher Scientific). Quality check was performed with the Bioanalyzer 2100 using the Agilent High Sensitivity DNA Kit (both Agilent, Santa Clara, USA). Pooled sequencing-ready libraries of the WTA and sample tag were sequenced on Illumina NovaSeq 6000 (150 bp, paired end reads; Eindhoven, The Netherlands) at Novogene Co Ltd (Beijing, China). Following service provider instructions, Seven Bridges Genomics (London, UK) was used to perform quality control, to map reads against mm10 using STAR and to determine expression counts. Data were further analysed using Seurat v.2.3Rpackage. The raw counts were normalized and log2 transformed by first calculating ‘size factors’ that represented the extent to which counts should be scaled in each library. Highly variable genes were detected using the proposed workflow of the scranR package and by applying false discovery rate≤0.05 and var.out$bio≥0.01 as cutoffs. Highly variable genes were subsequently used for unsupervised dimensionality reduction techniques and principal component analysis. Unsupervised clustering of the cells was performed using graph-based clustering based on SNN-Cliq and PhenoGraph as implemented in the Seurat v.2.3Rpackage (default parameters).

### Human subjects

Peripheral blood samples were obtained from HD and MS patients which were collected by Noorderhart hospital (Pelt, Belgium) and the University Biobank Limburg (UBiLim, Hasselt, Belgium) [69] after informed consent. Buffy coats of HD were purchased from the Belgian Red Cross. Samples were used immediately or stored in liquid nitrogen. Paired CSF and blood samples were collected from MS patients at diagnosis in the Klinikum Rechts der Isar, Technical University of Munich (Munich, Germany) after written informed consent. Donor characteristics are given in Table S3. Approval was obtained by the local ethical committees.

### Treg isolation

Tregs were isolated from peripheral blood mononuclear cells (PBMCs) by negative selection of CD4 (Miltenyi Biotec, Bergisch Gladbach, Germany) followed by positive CD25 selection (Miltenyi Biotec) using magnetic beads according to manufacturer’s instructions. Next, Tregs were sorted using a FACSAriaII or FACSAria Fusion (BD Biosciences) using following antibodies: CD4 APC-eFluor780, CD127 PE (all Thermo Fisher Scientific), CD25 PE/Cy7, CD25 Kiravia Blue 520, CD127 BV421, CD127 PE/Cy5, CD4 AF700 or CD4 BV421 (all Biolegend). The sorting strategy is depicted in Fig. S9 and purity of all sorts was confirmed to be >95%. Tregs (CD4^+^CD25^hi^CD127^lo^) and Teff (CD4^+^CD25^-^) were used in the Boyden chamber migration assay and the suppression assay.

### hCMEC/D3 cell culture

The human brain endothelial cell line hCMEC/D3 was obtained at Tébu-Bio (Le Parray-en-Yvelines, France). Cells were used up to passage 35. Cells were cultured in collagen coated (rat tail, type I, Sigma-Aldrich, Saint Louis, USA) culture flasks in growth medium (EGM-2 MV medium (Lonza, Basel, Switserland) supplemented with 2.5% fetal bovine serum (FBS, Gibco, Thermo Fisher Scientific)). Cells were collected using trypsin and cultured in 24 well plate or in Thincerts (both Greiner Bio-One) for migration assays as described further. After reaching near-confluency (90%), cells were replenished with experimental medium (basal EBM2 (Lonza) supplemented with Gentamicin (10 μg/ml, Sigma-Aldrich), Amphotericin B (1 μg/ml, Sigma-Aldrich), fibroblast growth factor (FGF, 1 ng/ml, Sigma-Aldrich), Hydrocortisone (HC, 1.4 μM, Sigma-Aldrich), 2.5% FBS (Gibco)). One day later, cells were treated with TNF-α (100 ng/ml) and IFN-γ (10 ng/ml, all Peprotech, London, UK) [70] for 24 hours in reduced medium (basal EBM2 (Lonza) supplemented with Gentamicin, Amphotericin, FGF, 0.25% FBS) [71].

### Boyden chamber migration assay

For migration assays, the Boyden chamber migration assay was used as described before [31, 71]. hCMEC/D3 were trypsinized and cultured in collagen-coated Thincerts (24 well, translucent, 3 μm, Greiner Bio-One) at a density of 25×10^3^ cells/cm². On day 3 and 5, cells were replenished with experimental medium as described before. Growth of the monolayer was followed using measurement of trans epithelial electrical resistance (TEER) using EVOM2 Epithelial Voltohmmeter (World precision instruments, Hertfordshire, UK) and reached a plateau phase on day 6. Then, cells were treated with TNF-α (100 ng/ml) and IFN-γ (10 ng/ml, all Peprotech) for 24 hours in reduced medium as described before. Before adding Tregs, the inserts were washed and put in a new plate with fresh reduced medium. (Pre-treated) Tregs (4×10^5^-1×10^6^) were allowed to migrate for 24 hours or were cultured in reduced medium as a control condition (untouched). Tregs from the well (migrated) and from the insert (non-migrated) were collected for RNA isolation, flow cytometry or further *in vitro* culturing. For RNA isolation, Tregs were pelleted and stored at -80°C in RTL lysis buffer (RNeasy micro kit, Qiagen, Hilden, Germany) with β-mercapto-ethanol (1:100). For flow cytometry, Tregs were washed in PBS before staining. For further *in vitro* culturing, Tregs were collected in culture medium as described below.

### Suppression assay

For all suppression assays, Teff were labelled with CellTrace Violet (5μM, Thermo Fisher Scientific) prior to culturing alone (2×10^4^ cells / well; referred to as 1:0; in U-bottomed 96 well plate) or together with Tregs. Cells were stimulated with MACS Treg Suppressor Inspector Beads (Miltenyi Biotec) in culture medium for 5 days at 37°C and 5% CO_2_. *Ex vivo* were immediately co-cultured after isolation. In the rescue experiments, Treg fractions were first treated with rapamycin (2 μM in DMSO, Sigma-Aldrich) or DMSO alone (control; 1/200, Sigma-Aldrich) for 4 hours in a U-bottom 96 well plate at a density of 6×10^5^ cells/well. Before addition to Teff or to the Boyden chamber migration assay, Tregs were washed in culture medium. Ratio’s at which cells were combined are specified in the appropriate figure legends. Gating strategy is depicted in Fig. S2. Proliferation controls were included and consisted of Teff cultured alone at the highest cell density. As a quality control, threshold for donor inclusion was set at minimal 70% proliferation for the 1:0 condition. Next, using flow cytometry, cell viability and CellTrace Violet dilution were analysed after 5 days. Prior to flow cytometry staining, culture supernatant was collected and stored at −80 °C.

### Bulk RNAseq

mRNA was isolated from Boyden chamber migration assay-derived Treg pellets from 5 HD and 4 uRRMS patients using RNeasy micro kit (Qiagen) according to manufacturer’s instructions. RNA concentration and quality were determined using a NanoDrop spectrophotometer (Isogen Life Science, IJsselstein, The Netherlands). Per sample, an amount of 1 ng of total RNA was used as input for the SMART-Seq® Stranded Kit (Cat. No. 634444; protocol version “022829”; low input option, Takara Bio USA, San Jose, USA). This kit can deal with degraded as well as high-integrity input RNA. Positive and negative controls were included in the experimental design using 1 ng Control RNA (included in the kit) and 7 μl RRI water, respectively. First, RNA is converted to cDNA using random priming (scN6 Primer) and SMART® (Switching Mechanism At 5’ end of RNA Template) technology and then full-length adapters for Illumina sequencing (including specific barcodes for dual-indexing libraries) are added through PCR using a limited number of cycles (5). The PCR products are purified and then ribosomal cDNA is selectively depleted by cleaving the ribosomal cDNAs by scZapR in the presence of mammalian-specific scR-Probes which target nuclear and mitochondrial rRNA sequences. This process leaves the library fragments originating from non-rRNA molecules untouched. The remaining cDNA fragments are further amplified with primers universal to all libraries (14 cycles). Lastly, the PCR products are purified once more to yield the final cDNA library. All libraries were finally quantified using Qubit dsDNA HS kit (Thermo Fisher Scientific) and their size distribution was checked using a Bioanalyzer 2100 (Agilent). Sequence-libraries of each sample were equimolarly pooled and sequenced on Illumina NovaSeq 6000 (100 bp, Single Reads, v1.5, 100-8-8-0) at the VIB Nucleomics Core (www.nucleomics.be, Leuven, Belgium). Quality of saw sequence reads was checked using FastQC version 0.11.8 and nucleotide calls with a quality score of 28 or higher were considered high quality. Adapters were removed using cutadapt v.2.4. Reads were aligned to the hg19 genome reference, using STAR (2.2.0e) and a maximum of five mismatches were allowed. Gene counts were retrieved using htseq-count using the “union” option. After removing absent features (zero counts in all samples), the raw counts were imported to R/Bioconductor package DESeq2 v.3.9 to identify differentially expressed genes among samples. The default DESeq2 options were used, including log fold change shrinkage. Differentially expressed genes were considered only significant when the Benjamini-Hochberg adjusted p-value (false discovery rate (FDR)) was <0.05. Heatmaps were created using the gplots::heatmap.2() function on transformed raw counts.

Ingenuity Pathway Analysis (IPA, Qiagen) was used to map lists of significant genes (FDR < 0.05) to gene ontology groups and biological pathways. The functional and canonical pathway analysis was used to identify the significant biological functions and pathways. Functions and pathways with p-values less than 0.05 (Fischer’s exact test) were considered statistically significant.

The gene set enrichment analysis (GSEA) was done using GSEA software, which uses predefined gene sets from the Molecular Signatures Database (MsigDB). For the present study, we used all the H: hallmark gene sets for GSEA analysis and a list of ranked genes based on a score calculated as log10 of p-value multiplied by sign of fold change. The minimum and maximum criteria for selection of gene sets from the collection were 10 and 500 genes, respectively [72].

### Bead-based immunoassay

hCMEC/D3 were trypsinized and cultured in collagen-coated 24 well plates (Greiner Bio-One) at a density of 25×10^3^ cells/cm². After reaching near-confluency (90%), cells were replenished with experimental medium as described before. The day after, cells were treated with TNF-α (100 ng/ml) and IFN-γ (10 ng/ml, all Peprotech) for 24 hours in reduced medium as described before. After 24 hours, hCMEC/D3 cells were washed once and put on fresh reduced medium. After another 24 hours, supernatant were collected and stored at −80 °C. To look at cytokine production by Tregs and Teff, the supernatants of suppression assays were collected after 5 days and stored at -80°C. Samples were thawed and used undiluted in LEGENDplex™ Human Anti-Virus Response Panel (Biolegend) according to manufacturer’s instructions to measure different cytokines. Samples were acquired using the LSRFortessa (BD Biosciences) and analysed using LEGENDplex™ Data Analysis Software (Biolegend). Concentrations were calculated by the use of the standard curves following the software guidelines.

### qPCR

To determine gene expression levels, hCMEC/D3 were trypsinized and cultured in 24 well plates (Greiner Bio-One) at a density of 25×10^3^ cells/cm². After reaching near-confluency (90%), cells were replenished with experimental medium as described before. The day after, cells were treated with TNF-α (100 ng/ml) and IFN-γ (10 ng/ml, all Peprotech) for 24 hours in reduced medium as described before. After 24 hours, hCMEC/D3 cells were washed once and put on fresh reduced medium. After another 24 hours, cell pellets were collected and stored at −80 °C. Cells were lysed using RLT lysis buffer (Qiagen) with β-mercapto-ethanol (1:100) and RNA was extracted using RNeasy micro kit (Qiagen), according to the manufacturer’s instructions. mRNA concentration was measured with Nanodrop (Thermo Fisher Scientific). Conversion to cDNA was performed using qScript cDNA supermix (Quanta Biosciences, Beverly, USA). Quantitative PCR was performed on a StepOne detection system (Thermo Fisher Scientific). Primer sequences are listed in the supplementary information (Table S4). Relative quantification of gene expression was accomplished using the comparative Ct method. Data were normalized to the most stable reference genes using geNorm.

### Flow cytometry

PBMCs were obtained via ficoll density gradient centrifugation (Cederlane lympholyte, Sheffield, UK) and frozen in liquid nitrogen for storage (10% DMSO in FBS (Gibco)). At the time of analysis, samples were thawed and rested for 4 hours at 37°C, 5% CO_2_ culture medium (RMPI-1640 (Lonza) supplemented with 10% FBS (Gibco), 1% nonessential amino acids, 1% sodium pyruvate, 50 U/ml penicillin and 50 U/ml streptomycin (all Life Technologies, Merelbeke, Belgium)). 26 HD, 24 uRRMS, 25 flRRMS and 23 uSPMS patients were included. Fresh paired CSF and blood samples of 5 uRRMS patients were used immediately for phenotyping. Treg fractions collected from the Boyden chamber assay were used directly for staining. After live/dead staining using Fixable Viability Dye eFluor 506 or LIVE/DEAD Fixable Aqua Dead Cell Stain Kit (Thermo Fisher Scientific), following antibodies were used: CD25 BB515, CCR8 BV650, RORγt R718 (all BD Biosciences), CD127 PerCP/Cy5.5, Helios PerCP/Cy5.5, FOXP3 BV421, CCR5 PE, CD11a AF597, CCR6 BV650, CCR6 PE/Cy7, CD49d BV605, CXCR3 BV711, MCAM APC, CD3 AF700, IL-6R PE/Cy7, CCR4 PE/Dazzle 594 (all Biolegend) and CD4 APC-eFluor780 (Thermo Fisher Scientific). Cells were permeabilized using the FOXP3/Transcription factor staining buffer kit (Thermo Fisher Scientific) according to manufacturer’s instructions. Human BD Fc block (BD Biosciences) was also used for paired blood-CSF staining. Samples were acquired using the LSRFortessa (BD Biosciences) and analysed using FlowJo 10.8.0 (BD Biosciences) and OMIQ (FlowSOM analysis, Santa Clara, USA). Percentages and median fluorescent intensity (MFI) are given.

### Statistics

Statistical analyses were performed using GraphPad Prism version 9.2.0 (GraphPad Software, San Diego, USA). Details of statistical tests are given in figure legends. Tests used are: 2way ANOVA with Bonferroni’s multiple comparisons test, (one-tailed) Wilcoxon test, Friedman test, Kruskal-Wallis with with Dunn’s multiple comparison test and the Mann-Whitney test. Cumulative data are shown as mean ± SEM. A (adjusted) p-value < 0.05 was considered significant.

## Authors’ contributions

Conceptualization: BB; methodology: PB, IH, CH, LA, MH; formal analysis: PB, IH, LA, MH; investigation: PB, LA, MH; writing-original draft: PB; writing-review and editing: BB, NH, TK, MK; visualization: PB, IH; supervision: MK, BB, NH. All authors have read and agreed to the published version of the manuscript.

## Acknowledgements

We thank our healthy donors for blood donations and Véronique Pousset and Anne Bogaers for assistance in collecting the blood samples. We thank the MS patients and team of nurses and neurologists from Noorderhart, Pelt, Belgium for the blood donation and collection. We thank the Biobank UBiLim for their service. We also thank Christel Bocken, Kim Ulenaers and Laura Dusaer for technical assistance. We thank Tebu-Bio for granting us to work with the hCMEC/D3 cell line. We thank Karsten Kretschmer, Hans Jörg Fehling and Adrian Liston for kindly providing us the FOXP3^Cre-GFP^ Rosa^RFP^ fate-mapping mice. We thank Nathalie Cools and Susan Schlenner for fruitful discussions.

## Supplementary Figures and Tables

**Fig. S1.**
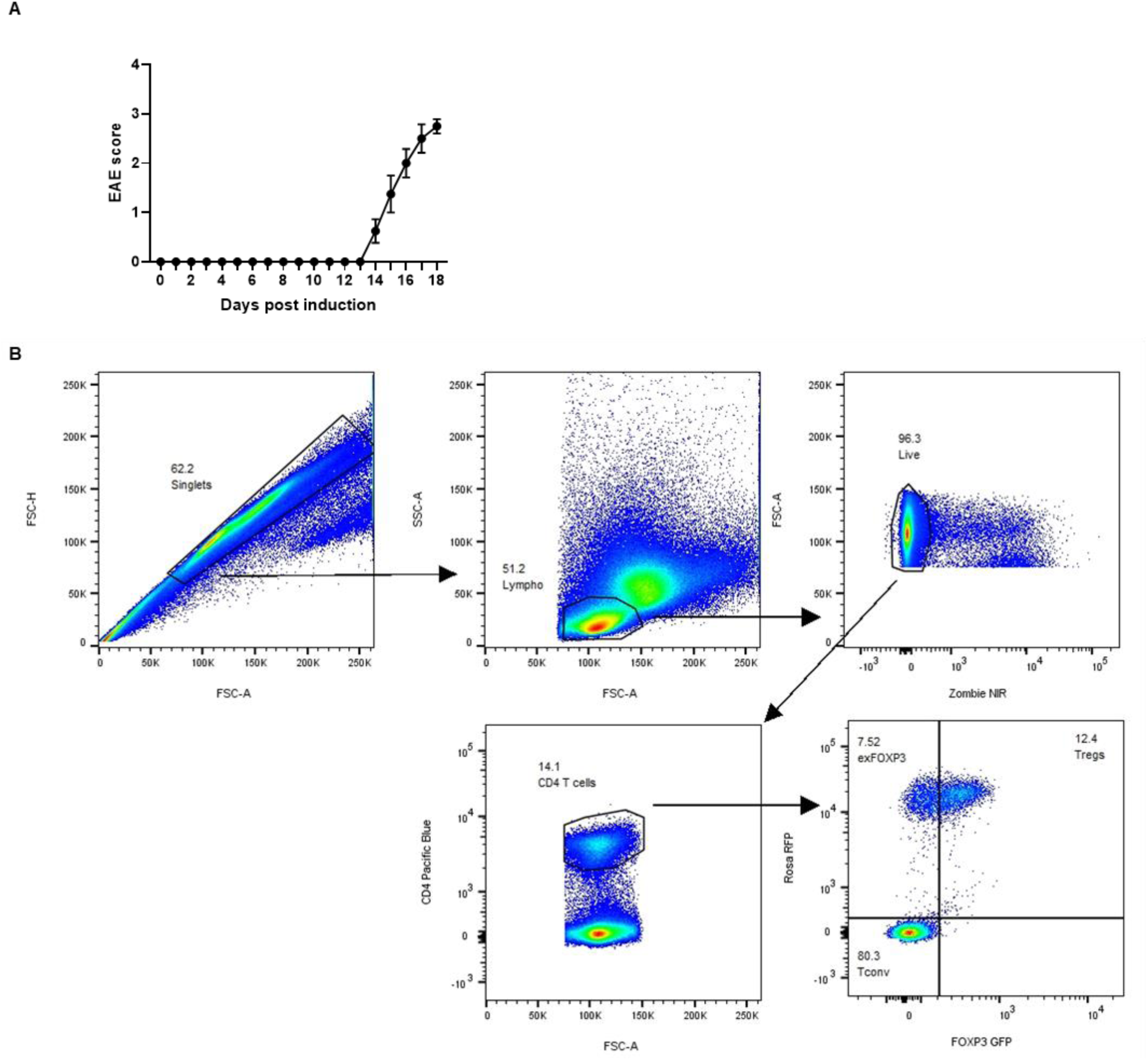
Disease development and gating strategy of FOXP3^+^ Tregs and exFOXP3 T cells *in vivo*. **A**. Course of EAE with onset (14 dpi) and peak (18 dpi). Animals and isolated cells were used for single-cell RNAseq. n=4. **B**. Using FOXP3^Cre-GFP^ Rosa^RFP^ fate-mapping mice, RFP^+^GFP^+^ FOXP3^+^ Tregs and RFP^+^GFP^-^ exFOXP3 T cells are studied in EAE. Gating strategy and representative plots of FOXP3^+^ Tregs and exFOXP3 T cells isolated from splenocytes are shown.

**Fig. S2.**
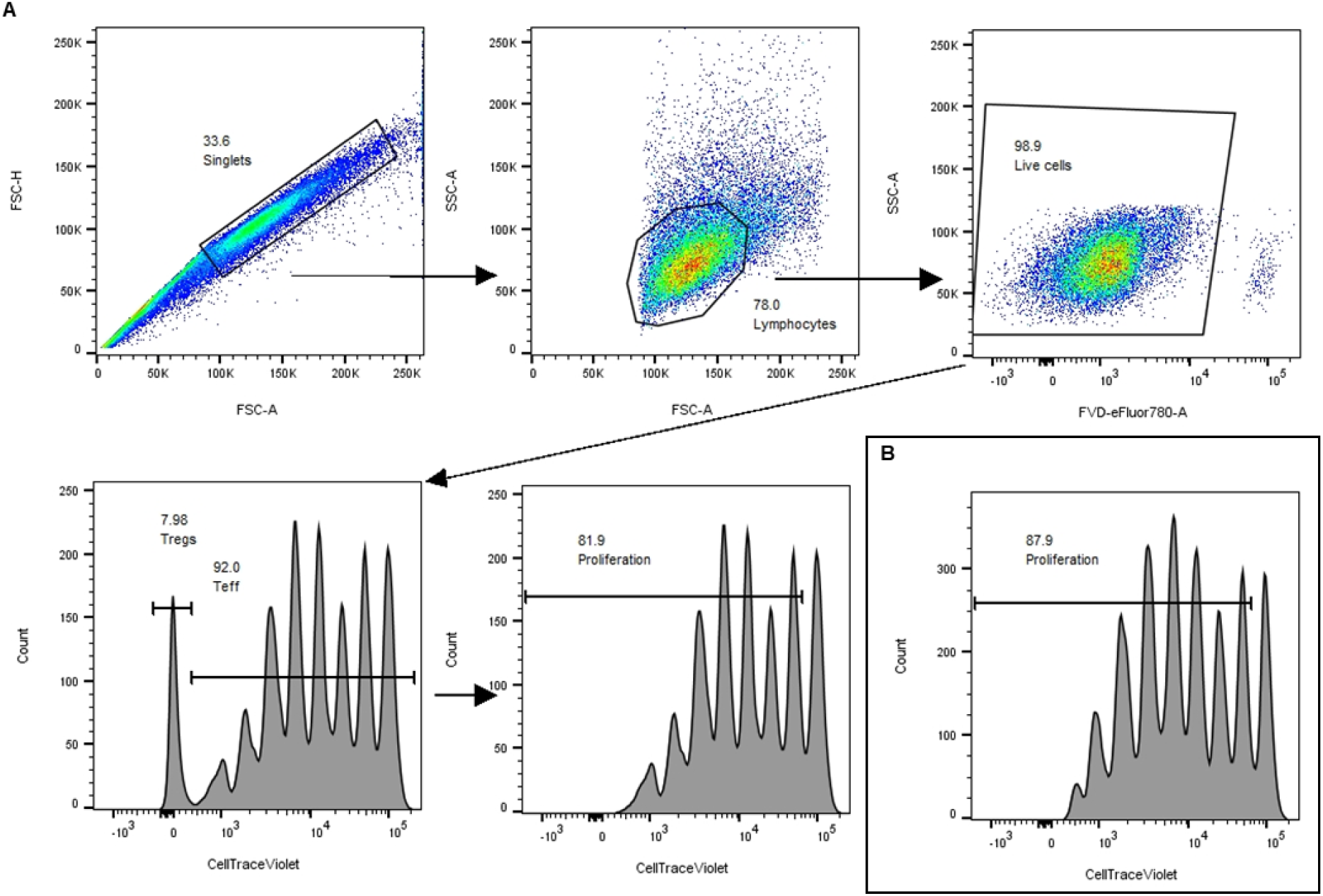
Gating strategy of suppression assay. Stimulated CellTrace Violet-labelled Teff are cultured for 5 days with and without Tregs. The dilution of the CellTrace is used as a measurement for proliferation. Gating strategy and representative plots of Teff proliferation cultured with migrated Tregs (A, 1:1 condition) or cultured alone (B, 1:0 condition).

**Table S1:** Pathway analysis of migrated vs. non-migrated Tregs using ingenuity pathway analysis (IPA).

| Top 10 IPA pathways | HD |  |  | uRRMS |  |  |
| --- | --- | --- | --- | --- | --- | --- |
|  | Rank | -log (p-value) | z-score | Rank | -log (p-value) | z-score |
| EIF2 Signaling | 1 | 37.5 | -6.274 | 1 | 41.9 | -6.14 |
| mTOR Signaling | 2 | 15.3 | -0.928 | 3 | 17.9 | -0.333 |
| Regulation of eIF4 and p70S6K Signaling | 3 | 15.3 | -1.606 | 4 | 16.4 | -0.853 |
| Coronavirus Pathogenesis Pathway | 4 | 14.9 | 4.695 | 2 | 18.9 | 4.529 |
| Th2 Pathway | 5 | 10.1 | 0.87 | 6 | 10.3 | 1.667 |
| Th1 and Th2 Activation Pathway | 6 | 9.67 | ND | 5 | 10.7 | ND |
| Molecular Mechanisms of Cancer | 7 | 8.81 | ND | 8 | 9.29 | ND |
| Senescence Pathway | 8 | 7.24 | 1.912 | 7 | 9.37 | 1.789 |
| RHOA Signaling | 9 | 7.17 | 2.466 |  | 4.74 | 2.401 |
| Protein Kinase A Signaling | 10 | 7.04 | -2.39 |  | 4.49 | -2.151 |
| ILK Signaling | 11 | 7 | -0.729 |  | 5.97 | 0.745 |
| PD-1, PD-L1 cancer immunotherapy pathway | 12 | 6.79 | -2.694 | 10 | 8.29 | -2.744 |
| IL-4 Signaling | 13 | 6.23 | ND |  | 4.73 | ND |
| Hepatic Fibrosis / Hepatic Stellate Cell Activation | 14 | 6.21 | ND |  | 6.2 | ND |
| Epithelial Adherens Junction Signaling | 15 | 5.82 | 1.508 | 14 | 7.68 | 1.697 |
| Th1 Pathway | 16 | 5.77 | 2.646 | 20 | 6.56 | 3.182 |
| Axonal Guidance Signaling | 17 | 5.68 | ND | 13 | 7.84 | ND |
| Actin Cytoskeleton Signaling | 18 | 5.62 | 2.714 | 18 | 6.83 | 3.434 |
| Pulmonary Fibrosis Idiopathic Signaling Pathway | 19 | 5.58 | 1.287 | 12 | 8.03 | 2.4 |
| ID1 Signaling Pathway | 20 | 5.56 | 0.555 |  | ND | ND |
| Role of Tissue Factor in Cancer |  | 4.64 | 0.287 | 9 | 8.33 | ND |
| Role of Macrophages, Fibroblasts and Endothelial Cells in Rheumatoid Arthritis |  | 3.72 | 0.212 | 11 | 8.24 | ND |
| Telomerase Signaling |  | ND | ND | 15 | 7.07 | 0 |
| Hepatic Fibrosis Signaling Pathway |  | ND | ND | 16 | 6.92 | 2.132 |
| Chronic Myeloid Leukemia Signaling |  | 3.37 | 0.264 | 17 | 6.83 | ND |
| p70S6K Signaling |  | 4.71 | 0.279 | 19 | 6.69 | -1.121 |
| Cell Cycle: G1/S Checkpoint Regulation |  | 3.6 | 0.289 | 20 | 6.59 | -1.091 |

ND: not determined

**Table S2:** Pathway analysis of migrated vs. non-migrated Tregs using gene set enrichment analysis (GSEA).

| Top 20 GSEA Hallmark pathways | HD |  |  | uRRMS |  |  |
| --- | --- | --- | --- | --- | --- | --- |
|  | Rank | NES | FDR | Rank | NES | FDR |
| Hallmark_Epithelial_Mesenchymal_Transition | 1 | 2.85 | 0.0001 | 1 | 2.69 | 0.0001 |
| Hallmark_Inflammatory_Response | 2 | 2.19 | 0.0001 | 12 | 1.95 | 0.0001 |
| Hallmark_MYC_Targets | 3 | -2.17 | 0.0001 | 9 | -2.10 | 0.0001 |
| Hallmark_G2M_Checkpoint | 4 | 2.14 | 0.0001 | 2 | 2.26 | 0.0001 |
| Hallmark_Mitotic_Spindle | 5 | 2.13 | 0.0001 | 6 | 2.19 | 0.0001 |
| Hallmark_Complement | 6 | 2.12 | 0.0001 | 11 | 2.01 | 0.0001 |
| Hallmark_Cholesterol_Homeostasis | 7 | 2.01 | 0.0001 | 8 | 2.17 | 0.0001 |
| Hallmark_P53_Pathway | 8 | 2.01 | 0.0001 | 12 | 1.97 | 0.0001 |
| Hallmark_Myogenesis | 9 | 2.01 | 0.0001 |  | 1.59 | 0.007 |
| Hallmark_TNFA_Signaling_via_NFKB | 10 | 1.99 | 0.0001 | 7 | 2.19 | 0.0001 |
| Hallmark_Apical_Junction | 11 | 1.98 | 0.0001 | 20 | 1.8 | 0.001 |
| Hallmark_E2F_Targets | 12 | 1.96 | 0.0001 |  | 1.73 | 0.002 |
| Hallmark_Reactive_Oxygen_Species_Pathway | 13 | 1.92 | 0.0001 | 3 | 2.25 | 0.0001 |
| Hallmark_Hypoxia | 14 | 1.90 | 0.0001 | 10 | 2.06 | 0.0001 |
| Hallmark_mTORC1_Signaling | 15 | 1.88 | 0.001 | 5 | 2.21 | 0.0001 |
| Hallmark_Apoptosis | 16 | 1.87 | 0.0001 | 4 | 2.24 | 0.0001 |
| Hallmark_Coagulation | 17 | 1.87 | 0.0001 |  | 1.36 | 0.039 |
| Hallmark_Allograft_Rejection | 18 | 1.84 | 0.001 | 15 | 1.91 | 0.0001 |
| Hallmark_Interferon_Gamma_Signaling | 19 | 1.83 | 0.001 |  | 1.75 | 0.002 |
| Hallmark_Kras_Signaling | 20 | 1.83 | 0.001 |  | 1.50 | 0.013 |
| Hallmark_Angiogenesis |  | 1.59 | 0.008 | 14 | 1.92 | 0.0001 |
| Hallmark_UV_Response |  | 1.8 | 0.001 | 16 | 1.89 | 0.0001 |
| Hallmark_IL6_JAK_STAT3_Signaling |  | 1.55 | 0.01 | 17 | 1.82 | 0.001 |
| Hallmark_IL2_STAT5_Signaling |  | 1.7 | 0.003 | 18 | 1.81 | 0.001 |
| Hallmark_PI3K_Akt_mTOR_Signaling |  | 1.37 | 0.047 | 19 | 1.8 | 0.001 |

NES: normalized enrichment score

**Fig. S3:**
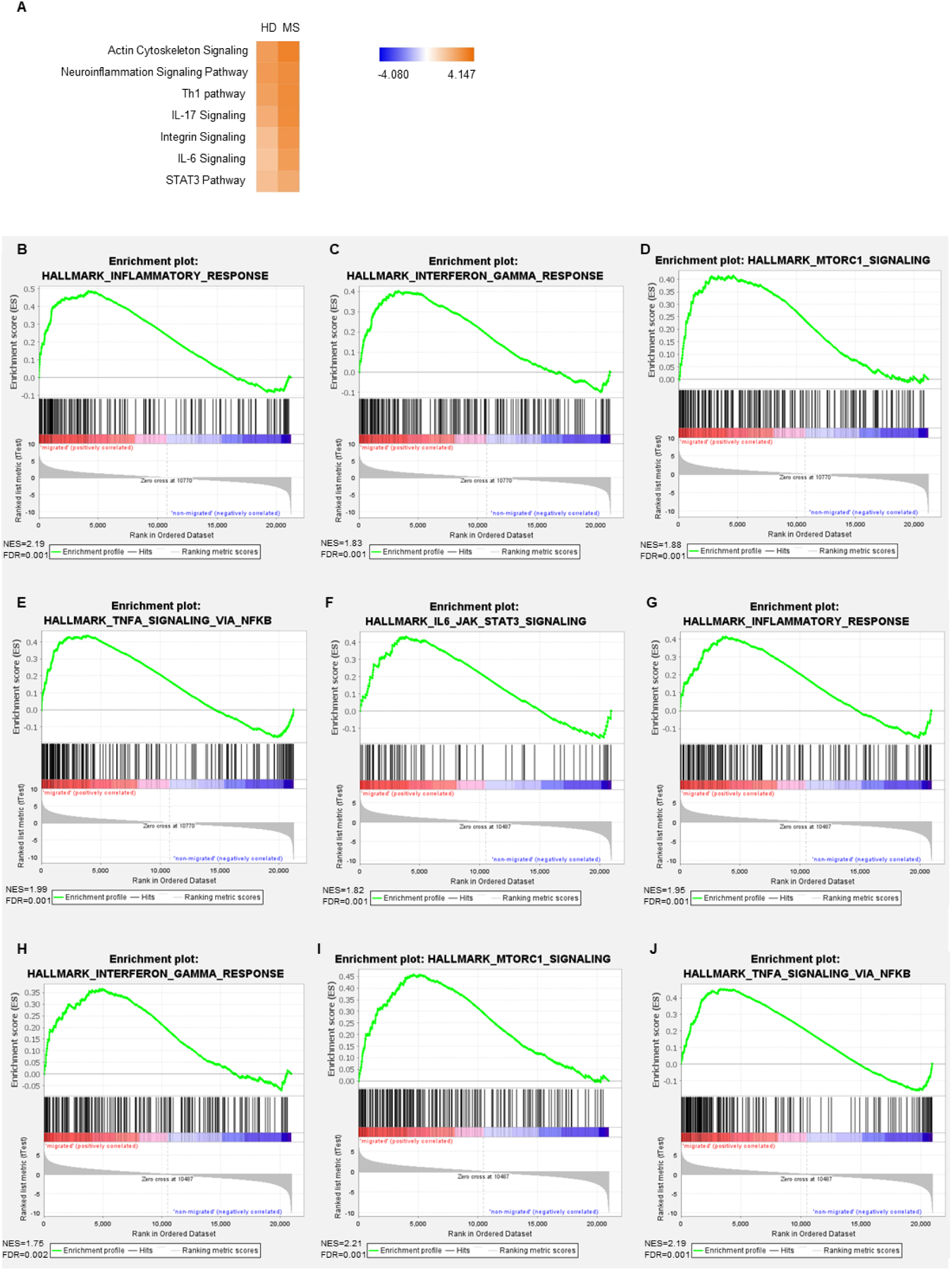
Pathway analysis using IPA and GSEA of transcriptomes of migrated vs. non-migrated Tregs isolated from HD and uRRMS patients. **A**. IPA predictions of canonical pathways on comparison analysis of migrated vs. non-migrated Tregs. Z-score predicts whether a canonical pathway is increased (positive z-score, orange) or decreased (negative, blue) in accordance with experimental datasets. Fischer’s exact test. **B-J**. GSEA enrichment plots (KEGG pathways) of inflammatory response, IFN-γ response, mTORC1 signaling, TNF-α signaling and IL-6-STAT3 signaling of migrated and non-migrated Tregs isolated from HD (**B-E**) and uRRMS patients (**F-J**). The green curve corresponds to the ES curve, which represents the running sum of the weighted enrichment score obtained from the GSEA algorithm. The red-blue band represents the degree of correlation of genes with the depicted pathway (red for positive and blue for negative correlation). The NES and corresponding FDR are reported within the plot. ES: enrichment score; KEGG: Kyoto encyclopedia of genes and genomes

**Fig. S4.**
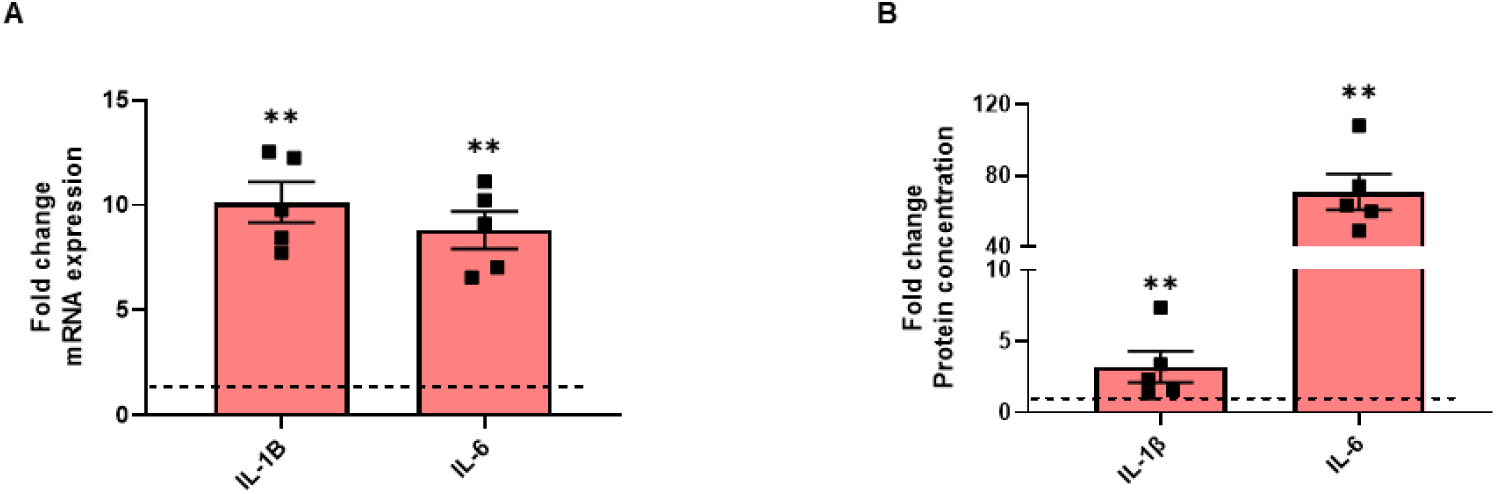
hCMEC/D3 cells produce IL-6 and IL-1β. hCMEC/D3 cells were left unstimulated or were TNF-α (100 ng/ml) and IFN-γ (10 ng/ml) stimulated for 24h. **A**. Fold change of *Il-1B* and *IL-6* mRNA expression shown compared to unstimulated cells (dotted line). n=5; Mann-Whitney test. **B**. Fold change of IL-1β and IL-6 concentration shown compared to unstimulated cells (dotted line). n=5; Mann-Whitney test. Data are represented as mean ± SEM. **: p<0.01.

**Fig. S5.**
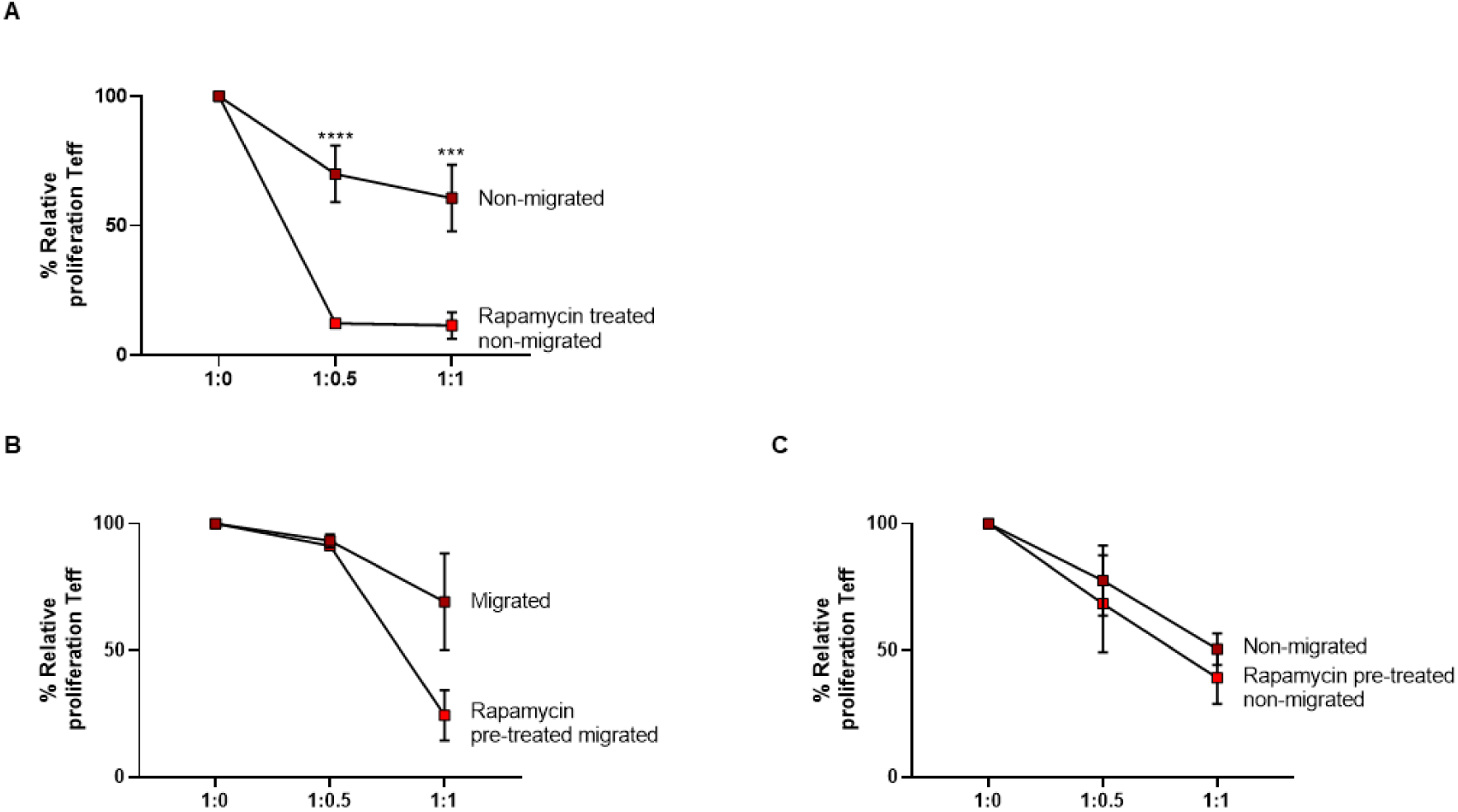
Rapamycin boosts suppressive capacity of non-migrated Tregs after migration but not before migration. **A**. Non-migrated HD-derived Tregs were collected and treated with rapamycin (2μM) for 4h, washed and co-cultured with Teff in a suppression assay. n=5. **B-C**. HD-derived Tregs were treated with rapamycin (2μM) for 4h, washed and loaded on the Boyden chamber migration assay. After 24h, cells were collected and co-cultured with Teff in a suppression assay. n=3. Percentage proliferation represents CellTrace dilution of Teff in suppression assays with different ratio’s of Teff and Tregs (given as Teff:Tregs). Relative proliferation is normalized to 1:0 condition (100%). Gating in Fig. S2.; 2way ANOVA with Bonferroni’s multiple comparisons test. Data are represented as mean ± SEM. ***: p<0.001; ****: p<0.0001.

**Fig. S6.**
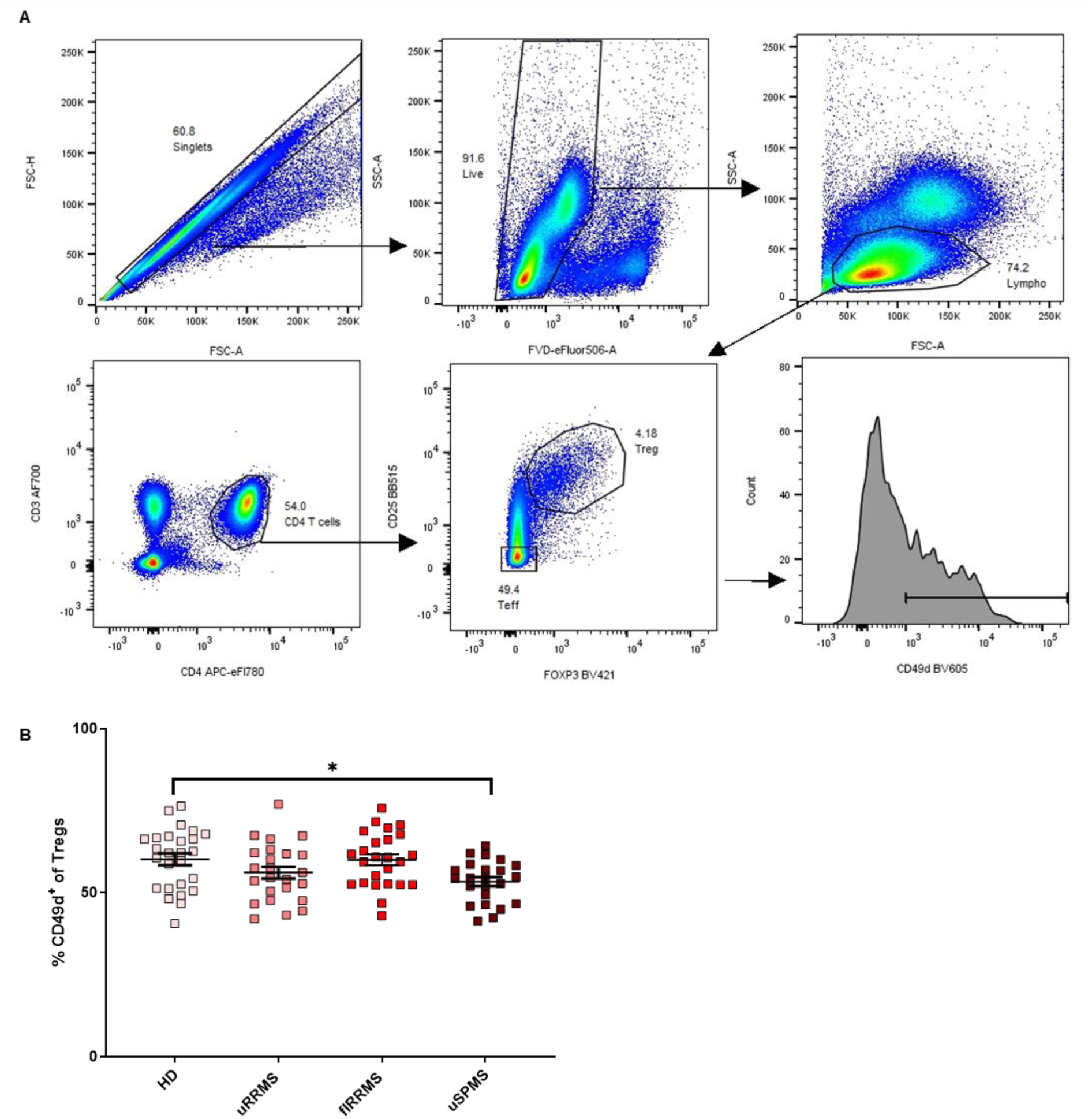
CD49d expression is not increased in MS patients. Frozen PBMC samples of HD, uRRMS, flRRMS and uSPMS patients are thawed and studied with flow cytometry. **A**. Gating strategy and representative plots of Tregs are shown. **B**. CD49d^+^ Treg not increased in differ between patient groups. Data are represented as mean ± SEM. n=26 (HD), n=24 (uRRMS), n=25 (flRRMS), n=23 (uSPMS); Kruskal-Wallis test with Dunn’s multiple comparisons test, *: p<0.05.

**Fig. S7.**
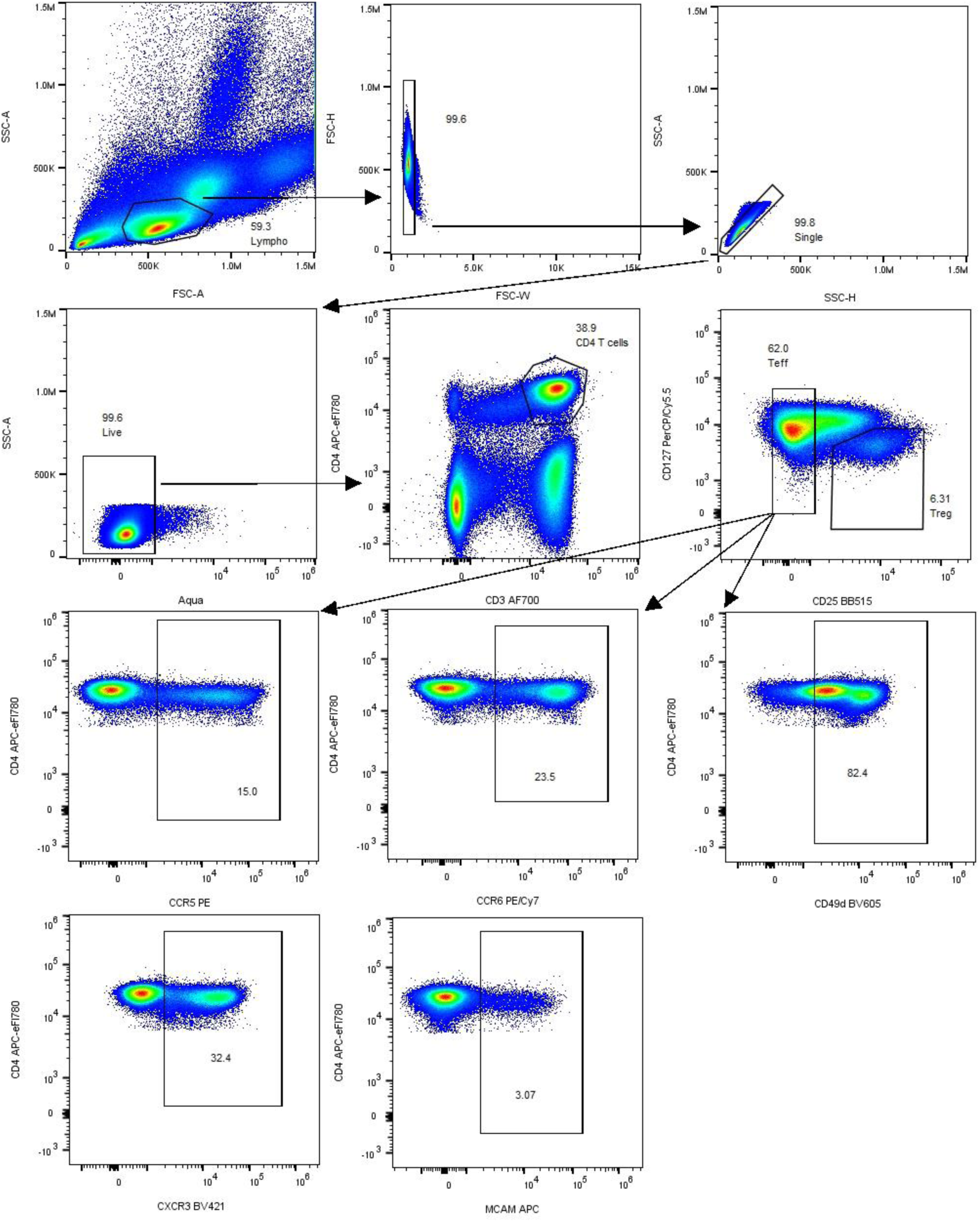
Gating strategy of migratory molecules on Tregs and Teff from paired blood and CSF samples of uRRMS patients. Fresh PBMCs and CSF samples are taken to diagnose MS and studied with flow cytometry. Gating strategy and representative plots of peripheral blood Teff are shown of MS patients.

**Fig. S8.**
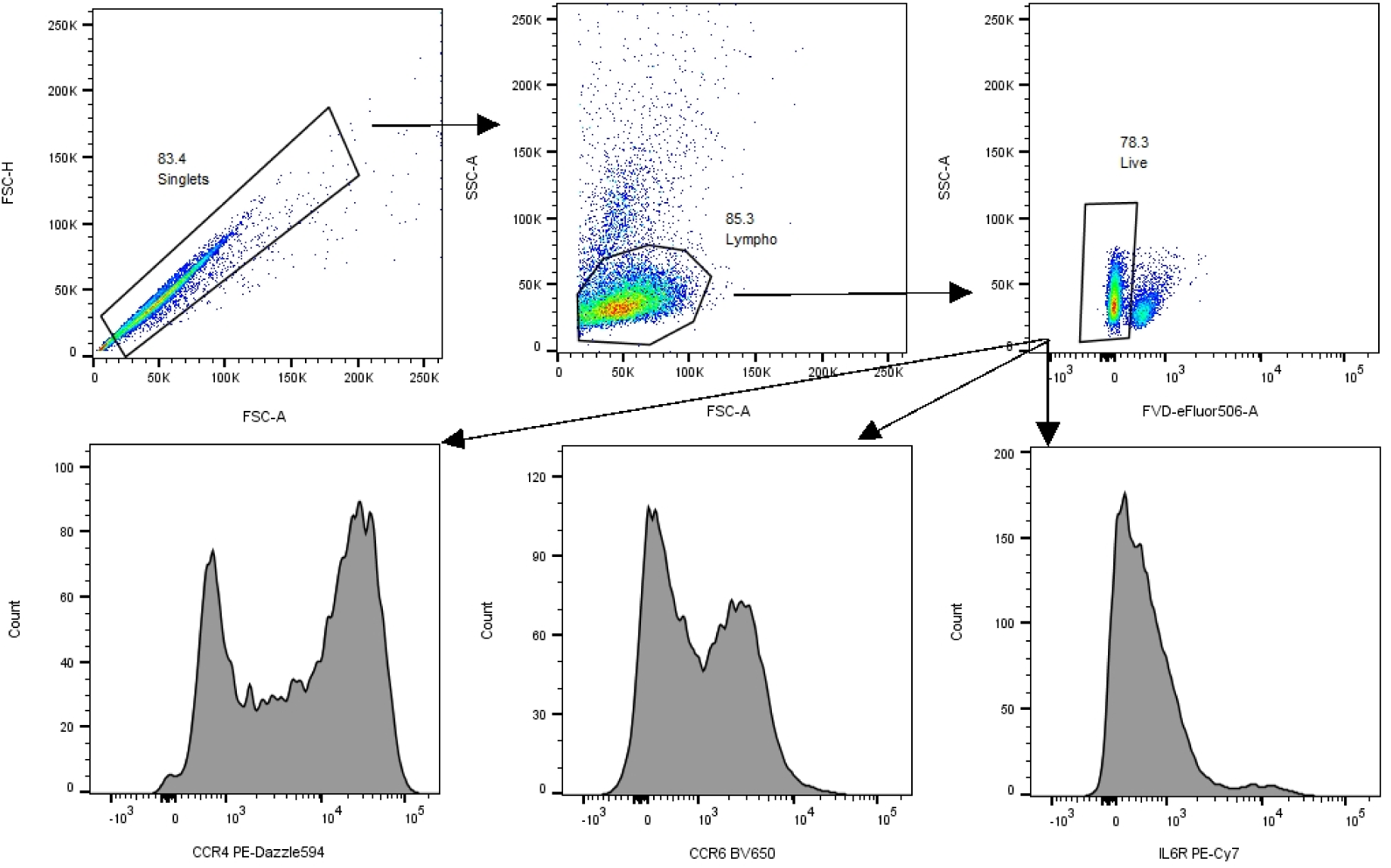
Gating strategy of RNAseq-identified hits on non-migrated and migrated Tregs using flow cytometry. Tregs are loaded on a Boyden chamber migration and the 2 cell fractions are stained. Gating strategy and representative plots of non-migrated Tregs are shown.

**Fig. S9.**
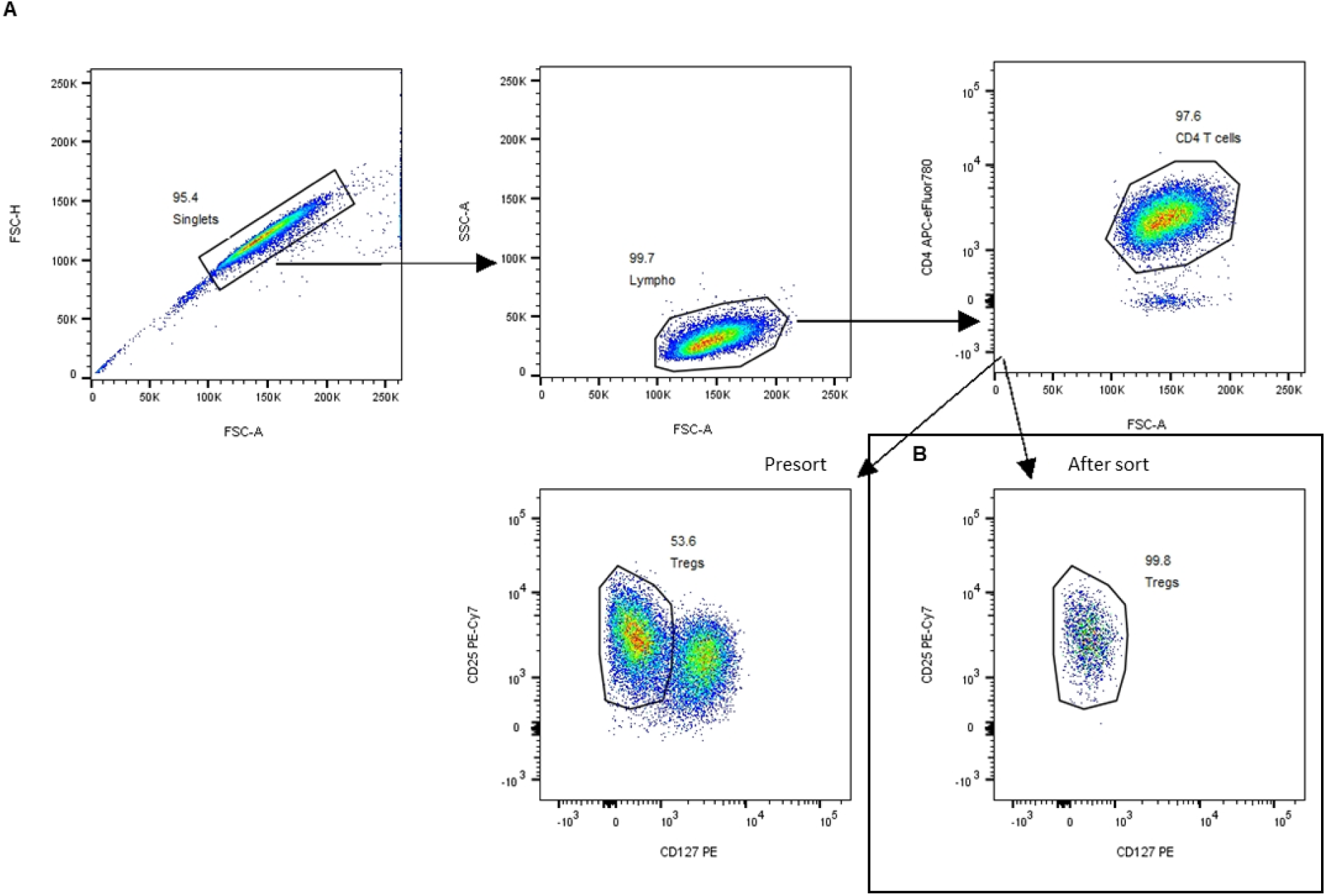
Gating strategy of human Treg sorting. **A**. Fresh PBMCs were sorted for Tregs based on CD4, CD25 and CD127 expression. **B**. Purity of sorted CD4^+^CD25^high^CD127^low^ Tregs.

**Table S3:**
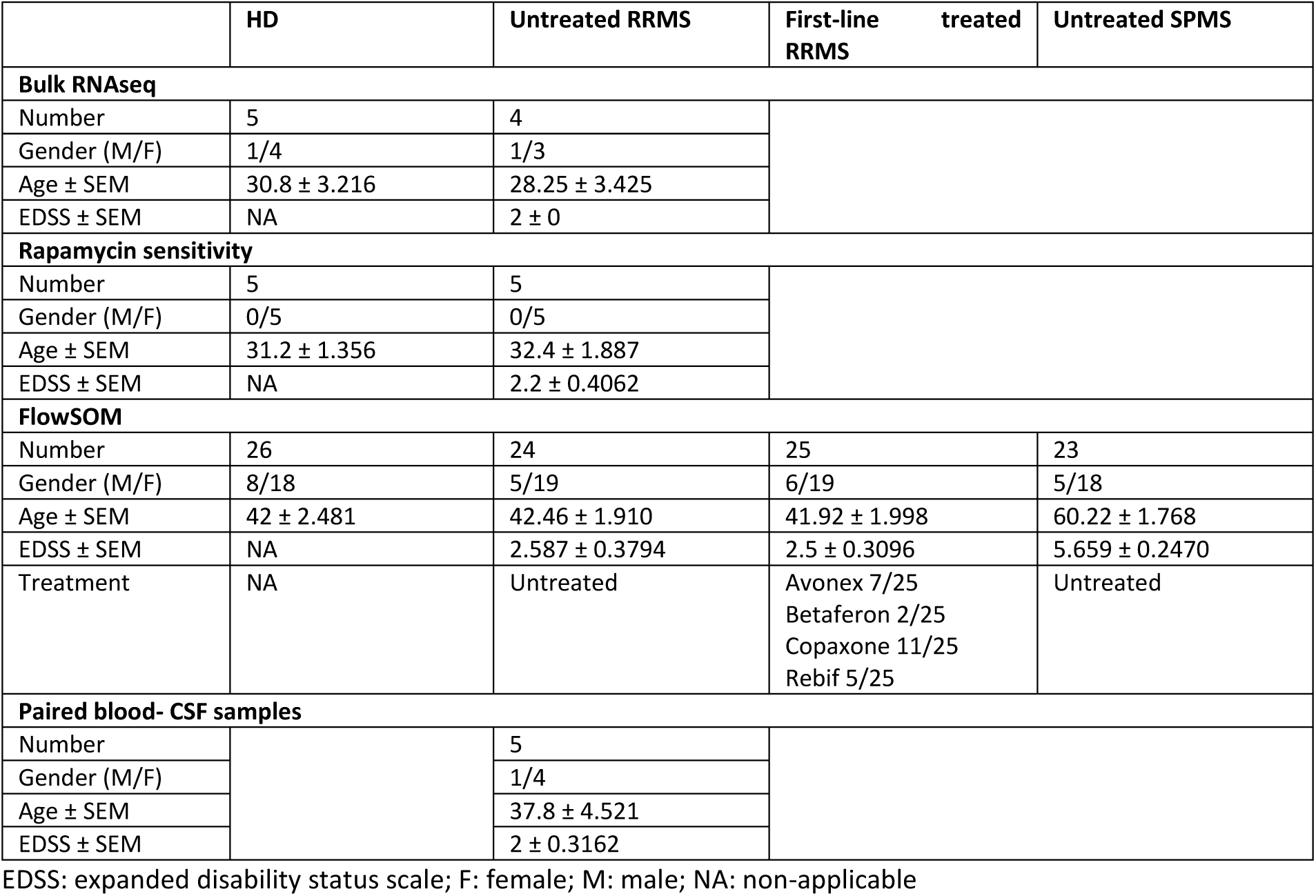
Patient information.

**Table S4:**
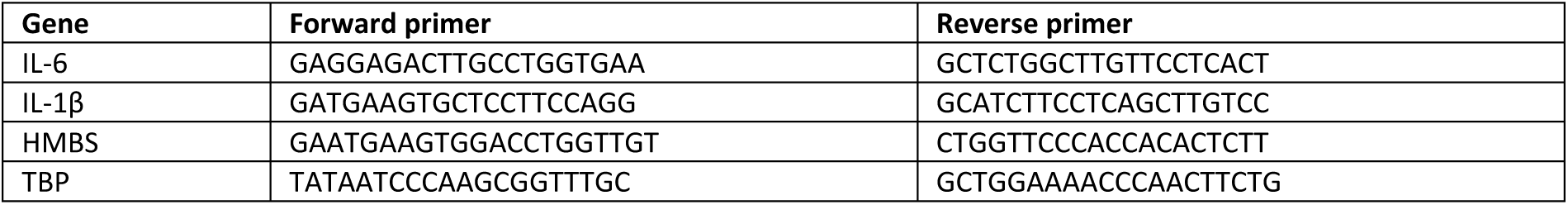
Primer sequences used for qPCR.

